# Genome-resolved metagenomics reveals conserved, flexible and emerging symbioses across global leafhoppers

**DOI:** 10.64898/2026.05.20.726650

**Authors:** Thierry Alexandre Pellegrinetti, Joshua Molligan, Abraão Almeida Santos, Nicolas Plante, Jordanne Jacques, Amélie Grégoire-Taillefer, Maria Cristina Canale, Maira Rodrigues Duffeck, Ashleigh M. Faris, Alejandro Olmedo-Velarde, Ivair Valmorbida, Edel Pérez-Lopez

**Author notes:** Corresponding author: EPL.

## Abstract

**Background:** Leafhoppers are among the most important insect vectors of plant pathogens worldwide and depend on microbial symbionts to exploit nutrient-poor phloem diets. However, most studies of leafhopper-associated microbiota have focused on a limited number of taxa or marker-gene surveys, leaving the genomic diversity, ecological organization, and functional potential of these microbial communities poorly understood. Here, we generated the Global Leafhopper Microbiome Catalog by integrating genome-resolved metagenomics from 171 leafhopper species across 11 subfamilies and 13 countries, including the first microbiomes characterized from Arctic leafhoppers.

**Results:** De novo assembly and genome reconstruction generated 337 high-quality non-redundant microbial genomes and 18.6 million non-redundant genes, substantially expanding the known microbial diversity associated with Cicadellidae, including several previously undescribed bacterial lineages. Comparative analyses revealed a recurrent modular microbiome architecture composed of: (i) a conserved core of obligate nutritional symbionts, dominated by ‘*Candidatus Karelsulcia’* and ‘*Candidatus Nasuia’*; (ii) a heterogeneous layer of secondary symbionts, including *Wolbachia*, *Arsenophonus*, *Rickettsia*, and *Diplorickettsia*; and (iii) a dynamic pool of environmentally acquired bacteria. While obligate symbionts remained highly conserved across divergent hosts, secondary and environmental taxa varied substantially among species and regions, suggesting repeated acquisition shaped by ecological filtering rather than host phylogeny alone. Comparative analyses between the specialist corn leafhopper *Dalbulus maidis* and the more polyphagous aster leafhopper *Macrosteles quadrilineatus* further showed that closely related vectors can maintain conserved ancestral symbionts while harboring markedly distinct accessory microbiomes. Arctic populations contained unique microbial assemblages enriched in functions associated with cold tolerance, oxidative stress, and reproductive manipulation. In addition, we identified numerous plant-associated bacteria, including phytoplasmas, spiroplasmas, *Pantoea*, and *Erwinia*, alongside taxa with predicted nutritional and plant growth-promoting functions.

**Conclusions:** Our findings reveal that leafhopper microbiomes are structured through the interaction of ancient obligate symbioses and flexible environmentally responsive microbial layers. This work establishes a genome-resolved framework for understanding microbiome evolution in insect vectors and highlights the potential role of microbial community structure in host adaptation, pathogen ecology, and sustainable pest management.

## INTRODUCTION

Leafhoppers (Hemiptera: Cicadellidae) are among the most ecologically and agriculturally important phloem-feeding insects and major vectors of plant pathogens^1–3^. Their success as herbivores and vectors relies not only on physiological adaptations but also on intimate associations with microorganisms that compensate for the nutritional limitations of a sap-based diet^4–6^. Most leafhopper species harbour the co-obligate endosymbionts ‘*Candidatus* Karelsulcia muelleri’ and the proteobacterial co-symbiont ‘*Candidatus* Nasuia deltocephalinicola’, which together synthesize essential amino acids^4,5^. These co-obligate symbionts are housed intracellularly within specialized bacteriocytes that form the bacteriome, a conserved symbiotic organ widely reported across hemipteran insects^4,10^. Additional facultative symbionts, including *Wolbachia*, *Rickettsia*, *Arsenophonus*, *Sodalis* and yeast-like fungi, can modulate host physiology, stress tolerance, and vector competence^5,7,8^. Leafhoppers transmit a wide diversity of plant pathogens, including bacteria such as mollicutes (phytoplasmas and spiroplasmas) and *Xylella fastidiosa*, as well as numerous plant viruses. Phloem-limited pathogens are primarily transmitted through circulative and often propagative mechanisms, whereas *X. fastidiosa* is transmitted in a non-circulative, foregut-borne manner ^9,10^. These interactions are highly specific and depend on molecular compatibility between insect, pathogen, and host plant, and emerging evidence suggests that resident microbial communities can influence vector competence and pathogen transmission^9^.

Given the dual roles of leafhoppers as hosts of complex microbial symbioses and as vectors of diverse plant pathogens, the diversity, evolutionary dynamics, and functional potential of leafhopper-associated microbiomes remain poorly characterized. Most studies have focused on a few taxa using microscopy or targeted molecular approaches, revealing remarkable variation in microbial communities among lineages such as Typhlocybinae and Eurymelinae and sensitivity to environmental factors, including host plant, temperature, and altitude^7,11,12^. However, systematic, genome-resolved surveys across host clades and eco-geographical contexts are still lacking, limiting our ability to understand symbiont turnover, metabolic specialization, and host adaptation.

In this study, we use large-scale genome-resolved metagenomics to provide a global view of symbiont diversity and functional potential in leafhopper microbiomes. We integrated 264 metagenomic datasets from 171 host species across 13 countries and clustered more than 78 million contigs at 95% identity, enabling gene-level analyses of metabolic and functional repertoires. This effort led to the reconstruction of 272 metagenome-assembled genomes, which, together with over 65 publicly available genomes, resulted in the creation of the Global Leafhopper Microbiome Catalog (GLMC), comprising 337 high-quality, non-redundant metagenome-assembled genomes (MAGs). The catalog incorporates newly generated data, including microbiomes from 38 original field samples representing 10 taxonomic groups collected along a latitudinal gradient from South America to the Canadian Arctic, capturing a broad range of ecological contexts. Collectively, these datasets establish a comprehensive genomic framework for leafhopper-associated microorganisms and provide a foundation for investigating the evolutionary plasticity of host–microbe interactions and their ecological and agricultural significance.

## RESULTS

### Gene-resolved catalog expands the microbial understanding of the leafhopper microbiome

To comprehensively profile leafhopper-associated microbiomes, we conducted a large-scale metagenomic analysis of 264 assemblies from 171 leafhopper species spanning 91 genera and 13 countries **(Figure 1a–b**, **Figure S1**, **Supplementary Table 1)**. This effort generated the most extensive genome-resolved dataset for the Auchenorrhyncha lineage within Hemiptera to date. Integration of these datasets resulted in the reconstruction of 78 million microbial contigs, which were gene-predicted and clustered into 18.6 million non-redundant open reading frames (ORFs), establishing the Global Leafhopper Microbiome Catalog (GLMC) **(Figure 1c**, **Supplementary Table 2)**. Approximately 51% (9.4 million) of these ORFs were taxonomically classified, with prokaryotic sequences accounting for 18.2% (3.4 million ORFs) of the total filtered for downstream analysis. Prokaryotic taxonomic profiling revealed that the GLMC was predominantly composed of Pseudomonadota (51.2%), Actinomycetota (15.1%), Bacillota (12.7%), and Bacteroidota (8.4%), with Enterobacterales, Burkholderiales, Pseudomonadales, and Bacillales identified as the most prevalent orders (**Figure 1d**, **Supplementary Table 3**).

**Fig. 1.**
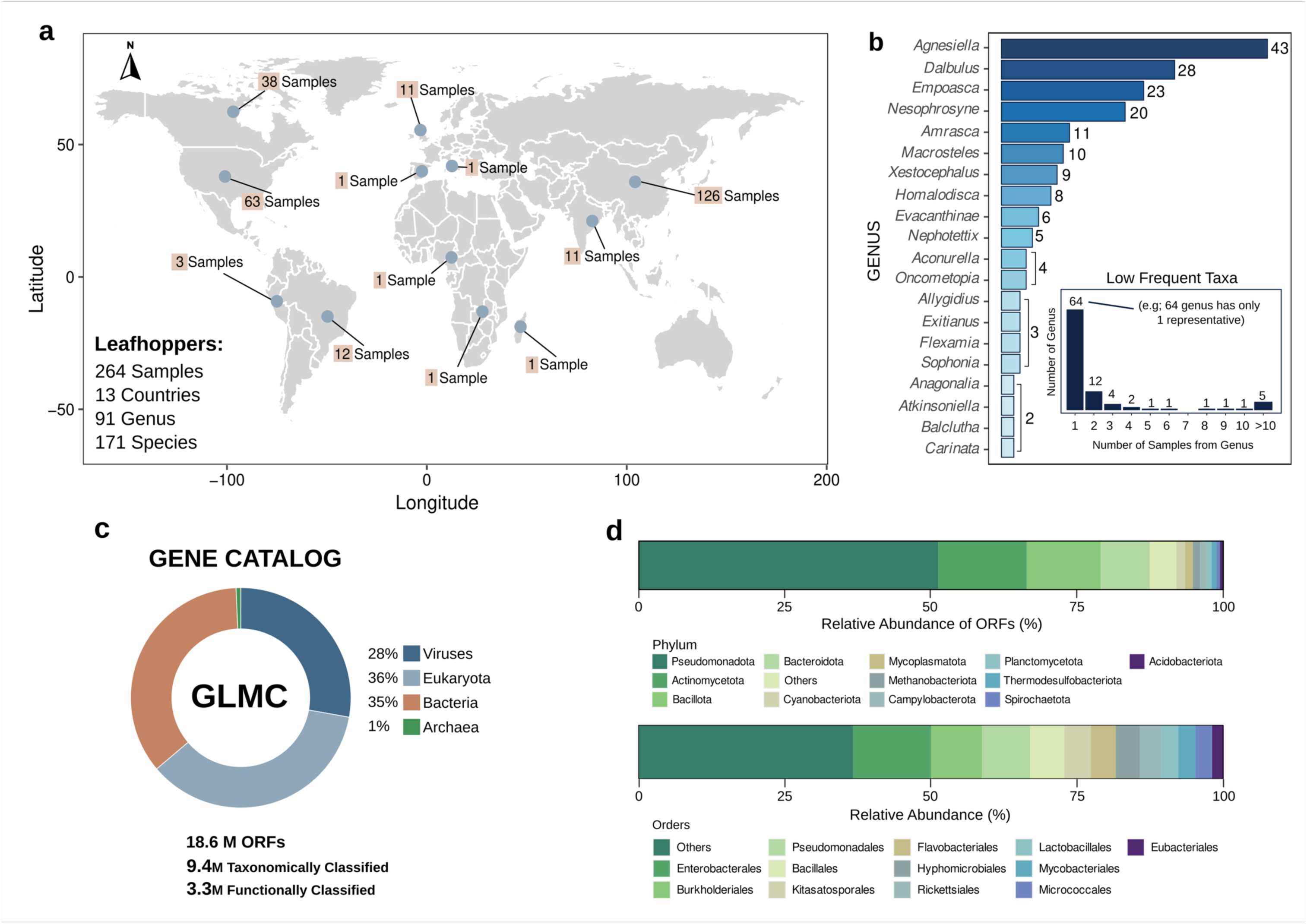
Global distribution and genomic landscape of leafhopper-associated microbiomes. **a**, Geographic distribution of the 264 leafhopper metagenomes analyzed in this study, spanning 13 countries and six continents [e.g., the Americas, Africa, Europe, and Asia]. **b**, Taxonomic diversity of the host dataset, comprising 171 leafhopper species across 91 genera and 11 subfamilies. The box inside the graph represents low represented genera including the number of genera presenting one to more than ten representatives (i.e. in the dataset, 64 genera have only one representative, 12 genus have 2 representatives and 3 genera have 4 representatives). **c**, General taxonomic classification metrics of Global Leafhopper Microbiome Catalog (GLMC), illustrating the accumulation of 18.6 million non-redundant open reading frames (ORFs) from 78 million reconstructed microbial contigs. **d**, Phylogenetic composition of the microbiome at the phylum and order level, highlighting the dominance of Pseudomonadota, Actinomycetota, and Bacillota as phyla, Enterobacterales, Burkholderiales, and Pseudomonadales as the top taxa across the surveyed leafhopper populations.

We successfully recovered 6,944 ribosomal genes, including 2,525 16S rRNA sequences, which were clustered at 97% identity and filtered by size to yield 278 OTUs spanning 59 bacterial genera **(Figure 2a**, **Supplementary Table 4)**. These findings indicate that leafhopper microbiomes are dominated by a limited number of core symbiont lineages, however, exhibit greater taxonomic breadth than previously reported^5,6,8,11,13^. Community structure was strongly species-dependent, with finer host taxonomic resolution corresponding to increasingly distinct microbial assemblages **(Supplementary Figure 2)**. Among the taxa identified, ‘*Candidatus* Karelsulcia’ and ‘*Ca*. Nasuia’ (hereafter Karelsulcia and Nasuia) were the most abundant **(Figure 2a–b)**, with Nasuia exhibiting the highest number of OTU variants, suggesting significant intraspecific diversification. Mapping of 16S rRNA reads across the metagenomic datasets revealed that putative symbionts and environmentally associated taxa, including *Wolbachia*, *Rickettsia*, *Pectobacterium*, *Arsenophonus*, *Spiroplasma*, and *Ca*. Baumannia were among the most prevalent taxa **(Figure 2b)**. Although ORF-level taxonomic assignment revealed a partially distinct pattern, Karelsulcia remained the most represented lineage, followed by *Wolbachia*, *Pseudomonas*, *Arsenophonus*, ‘*Ca*. Phytoplasma’, ‘*Ca*. Tisiphia’, and ‘*Ca*. Palibaumannia’ **(Figure 2c)**.

**Fig. 2.**
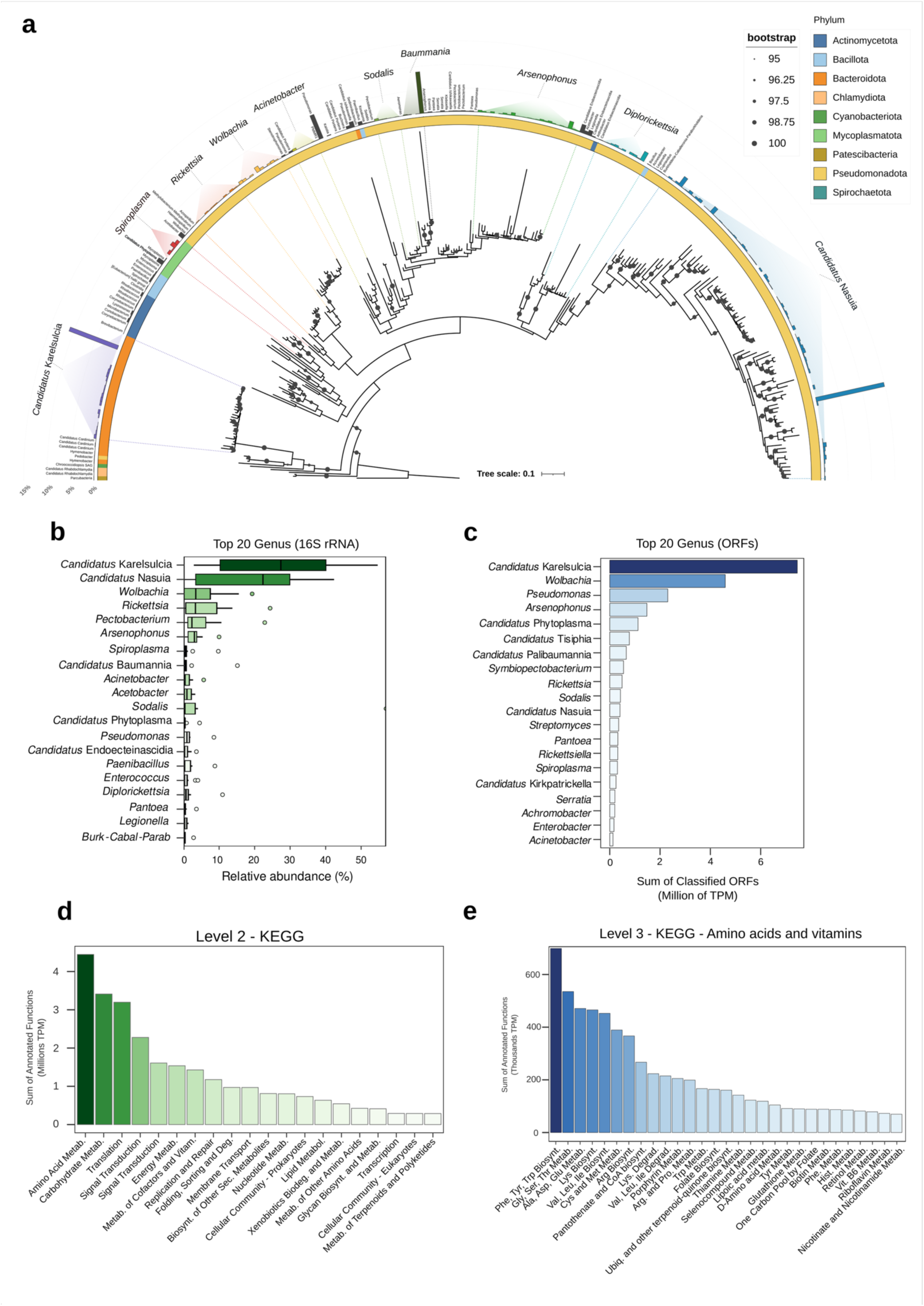
Phylogeny, taxonomic composition and metabolic functional potential of the leafhopper microbiome. **a**, Maximum-likelihood phylogenetic tree of 278 bacterial OTUs (97% identity) recovered from *de novo* assembly of 264 leafhopper metagenomes. Leaf nodes are color-coded by phylum, and the outer ring indicates the name of the genus. **b**, Relative abundance of the 20 most prevalent bacterial genera determined by mapping metagenomic reads to reconstructed 16S rRNA sequences. **c**, Taxonomic profile of the microbiome based on ORF-level coverage, showing the top 20 genera ranked by total recruited reads. Discrepancies between **b** and **c** reflect differences in copy number and resolution between marker gene and whole-genome approaches. **d**, Broad functional characterization of the leafhopper microbiome annotated at KEGG Level 2, displaying the relative abundance of major gene categories. **e**, Fine-scale functional analyses of metabolic pathways (KEGG Level 3) specifically focused on amino acid and vitamin metabolism. The bar plot highlights the metabolic enrichment of pathways involved in the biosynthesis of essential amino acids and B-vitamins (for example, thiamine, riboflavin and folate), underscoring the nutritional role of the bacterial symbionts.

Functional annotation of the taxonomically classified ORFs yielded 1.43 million assignments (41.98%) **(Supplementary Table 5)**. Leveraging ORF-to-read mapping, we observed a strong enrichment of pathways essential to leafhopper biology, including amino acid and carbohydrate metabolism, translation, signal transduction, energy production, and the biosynthesis of vitamins and cofactors **(Figure 2d, Supplementary Table 6)**. At KEGG level 3, these annotations indicate that aromatic amino acid biosynthesis and B-vitamin-related functions, specifically pantothenate, folate, and thiamine metabolism, represent the primary metabolic contributions provided by the leafhopper-associated microbiome **(Figure 2e**, **Supplementary Table 6)**.

### Leafhopper microbiomes are rich in uncharacterized microbial lineages

Using genome-resolved metagenomics, we reconstructed 272 non-redundant metagenome-assembled genomes (nrMAGs) from 112 leafhopper species across 64 genera, comprising 49.6% high-quality and 50.4% medium-quality genomes according to MIMAG standards **(Figure 3a**, **Supplementary Table 7)**. The nrMAGs exhibited high genome integrity, with a mean completeness of 86.8%, low contamination (0.64%), an average contig N50 of 124 kb, and a mean of 154 contigs per genome **(Figure 3b)**.

**Fig. 3.**
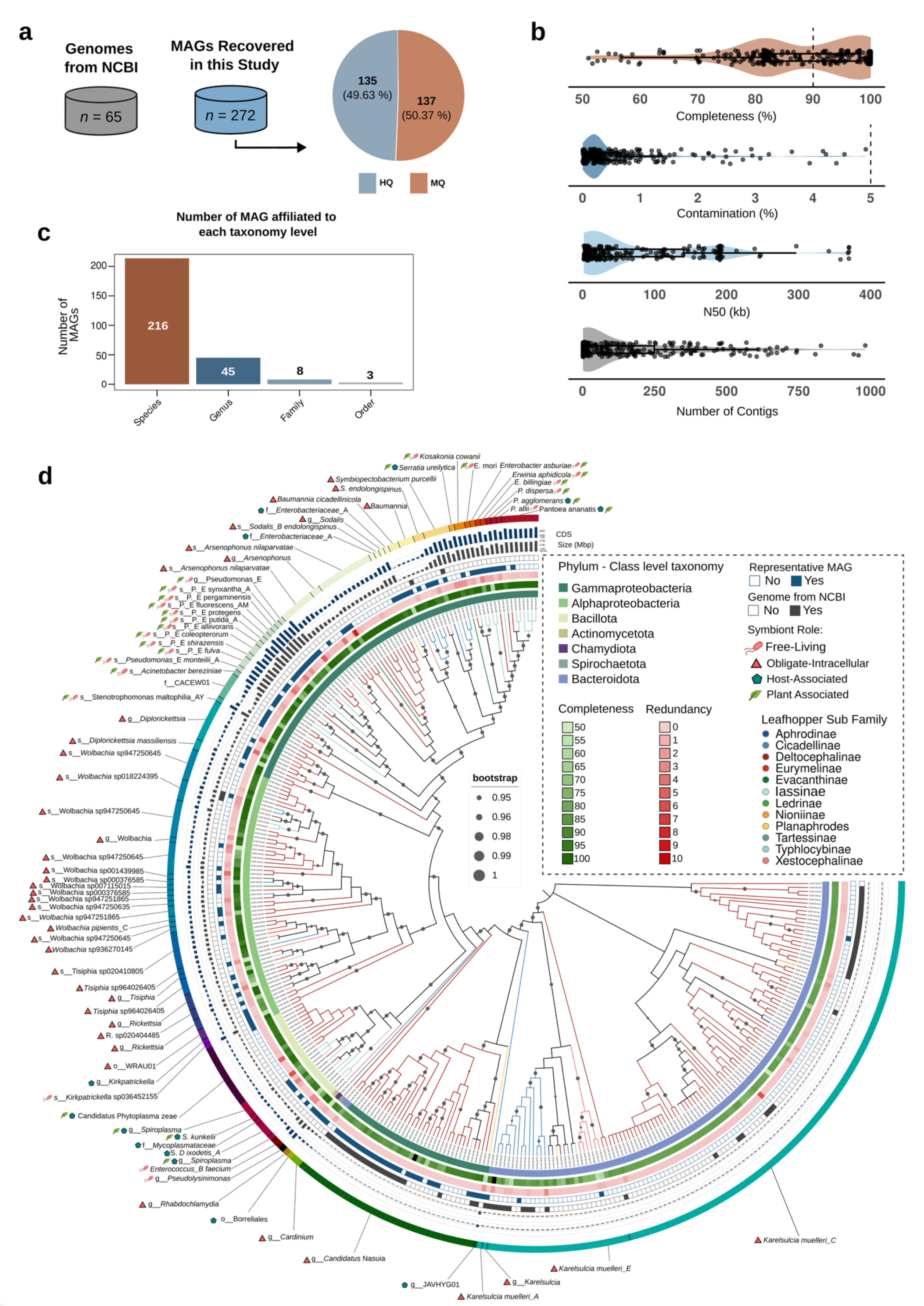
Genomic quality, taxonomic resolution and phylogenomic diversity of the extended leafhopper microbiome catalogue. **a**, Source and quality distribution of the analyzed genomes. The bar chart compares the number of publicly available genomes retrieved from NCBI (n=65, grey) with the novel metagenome-assembled genomes (MAGs) recovered in this study (n=272). The inset pie chart details the quality assessment of the novel MAGs, classified as high-quality (n=135) or medium-quality (n=137) according to MIMAG standards. **b**, Assessment of genomic integrity. Box plots overlaid with jitter points illustrate the distribution of key assembly metrics: completeness, contamination, contig N50 and total number of contigs per genome. Centre lines represent the median; box limits indicate the 25th and 75th percentiles. **c**, Taxonomic resolution of the catalogue. Bar plot displaying the count of MAGs successfully classified at the species (n=216), genus (n=45), family (n=8) and order (n=3) levels. **d**, Phylogenomic reconstruction and trait landscape of leafhopper-associated bacteria. Maximum-likelihood tree based on concatenated marker genes including all 337 analyzed genomes. Branch colors correspond to the host leafhopper subfamily, and nodes indicate bootstrap support values. Concentric rings (from inner to outer) denote: (1) bacterial phylum assignment; (2) genome completeness; (3) redundancy status; (4) representative status within the catalogue; and (5) data source (NCBI vs. this study). Outer bars represent genome size (Mbp) and coding sequence (CDS) count. Tip labels indicate bacterial species names, accompanied by symbols denoting the predicted lifestyle: free-living, obligate-intracellular, host-associated or plant-associated.

Taxonomic assignment resolved 216 nrMAGs to species level, 45 to genus level, eight to family level, and three to order level **(Figure 3c)**. After incorporating 65 publicly available genomes, the final dataset comprised 337 genomes covering eleven leafhopper subfamilies, dominated by Deltocephalinae (227), followed by Typhlocybinae (41), Cicadellinae (30), Xestocephalinae (12), and other subfamilies (27 genomes) **(Supplementary Table 8)**. Microbial genomes were recovered from 80 leafhopper species lacking any previously available symbiont genomes. Among the assembled genomes, the most prevalent symbionts were Karelsulcia (120 genomes) and Nasuia (47 genomes), consistent with their conserved co-obligate roles in provisioning essential amino acids in sap-feeding Auchenorrhyncha. A diverse set of putative secondary and facultative symbionts, as well as other leafhopper-associated bacteria, was also reconstructed, including *Wolbachia* (35 genomes), *Arsenophonus* (20 genomes), *Tisiphia* (12 genomes), *Diplorickettsia* (11 genomes), *Rickettsia* (7 genomes), and *Symbiopectobacterium* (7 genomes), expanding the available genomic resources for leafhoppers symbionts and leafhoppers-associated microbiota **(Figure 3d**, **Supplementary Tables 7 and 8)**.

Approximately 23% of the nrMAGs (43 genomes) lacked close relatives in the Genome Taxonomy Database (GTDB) r226, indicating the presence of poorly characterized or previously unclassified bacterial taxa **(Figure 4**, **Supplementary Table 9)**. Due to ANI (Average Nucleotide Identity) values below species thresholds, only 44 genomes could be assigned to the genus level. These include lineages affiliated within *Diplorickettsia*, *Rickettsia*, *Spiroplasma*, *Baumannia*, *Arsenophonus*, *Cardinium*, *Sodalis*, and *Wolbachia* **(Figure 4a**, **Supplementary Table 9)**. Among those genomes without conclusive taxonomic delineation, six were assigned to the potentially new family CACEW01 (here proposed as ‘*Candidatus* Edelium’), forming two distinct clades within the order Thiotrichales **(Figure 4b)**. These six genomes were consistently detected across multiple leafhopper hosts, including *Dalbulus maidis*, *Nesophrosyne* spp., and *Amrasca biguttula*. Nine nrMAGs assigned to *Diplorickettsia* exhibited low ANI values relative to *Diplorickettsia massiliensis* (84.3% to 88.2%) and formed a well-supported, distinct phylogenomic clade separate from other *Rickettsiella* and *Aquirickettsiella* lineages **(Figure 4c)**. These genomes were detected across multiple Cicadellidae genera, representing the first genomic characterization of a *Diplorickettsia*-like lineage associated with leafhoppers. In addition, based on ANI values, two nrMAGs were assigned to the potentially new family WRAU01 (here proposed as ‘*Candidatus* Diplorickettsia cicadellinicola’), forming a distinct clade adjacent to Holosporales and ‘*Candidatus* Pelagibacterales’ **(Figure 4d)**. These genomes were consistently recovered from the major crop pests *D. maidis* and *Graminella nigrifrons*. In addition to two potential new bacterial families, the genomes within these lineages could be proposed as representing up to five new species-level lineages. Further studies will be needed to characterize their biology and ecological relationships (**Supplementary Table 7**).

**Fig. 4.**
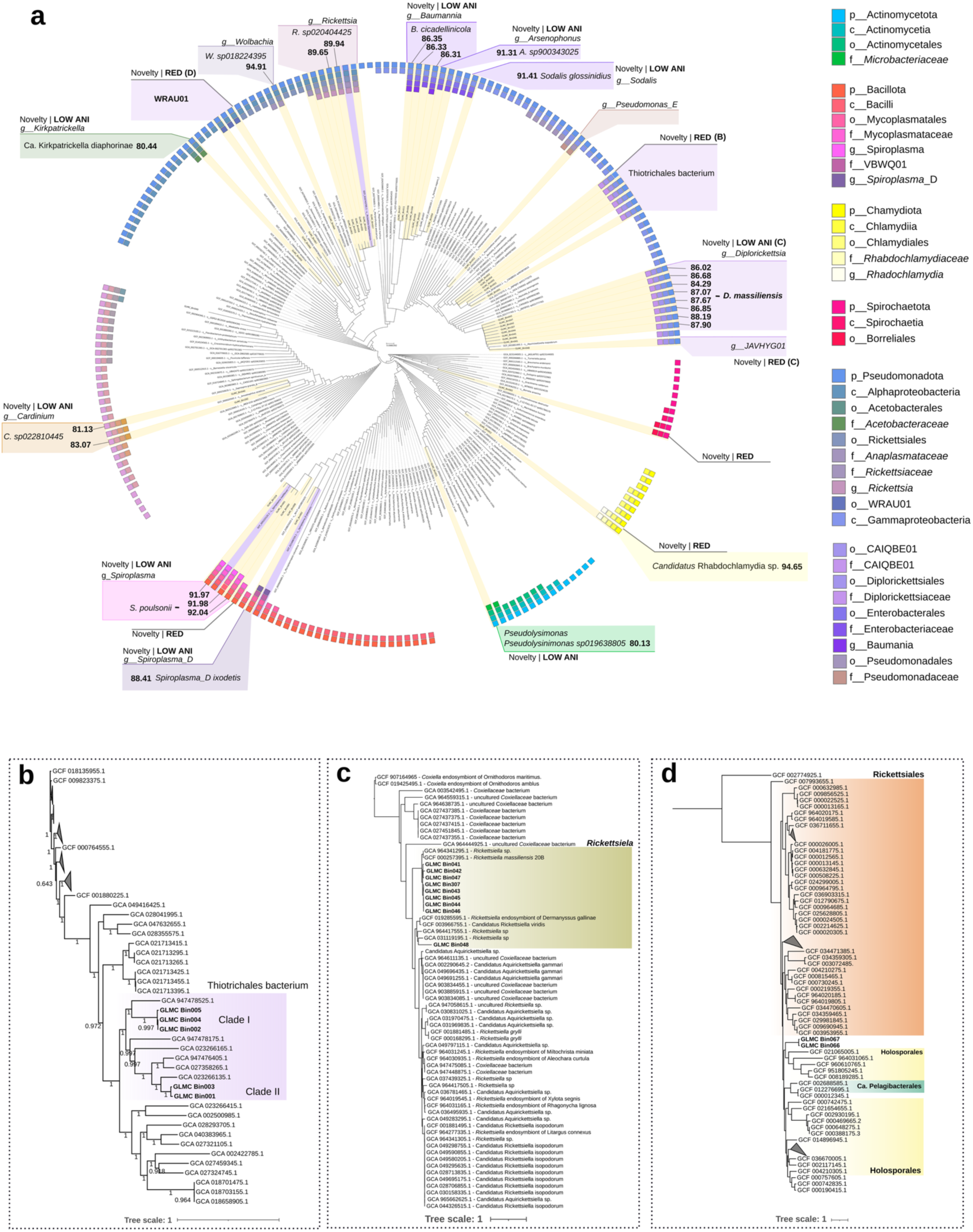
Phylogenomic placement and characterization of novel bacterial lineages within the leafhopper microbiome. **a**, Global phylogeny of unclassified taxa. Maximum-likelihood tree based on GTDB-Tk marker genes displaying metagenome-assembled genomes (MAGs) that could not be assigned to a known species (ANI <95%). These genomes represent putative novel taxa based on Average Nucleotide Identity (ANI) and Relative Evolutionary Divergence (RED) values. Outer rings denote the closest taxonomic assignment (family or genus level), while the associated colored box visualizes pairwise ANI values against the closest reference genomes. **b**, Phylogeny of the novel *Thiotrichales*-like lineage (proposed ‘*Candidatus* Edelium’). The tree resolves five MAGs assigned only to the order level into two distinct subclades: Clade I (GLMC_Bin002, Bin004 and Bin005) and Clade II (GLMC_Bin001 and Bin003). Both clades form a monophyletic group distinct from known aquatic *Thiotrichales* references, supported by low genomic similarity. **c**, Phylogenetic resolution of the *Diplorickettsia*– *Rickettsiella* complex (proposed ‘*Ca.* Diplorickettsia cicadellinicola’). The recovered MAGs form a cohesive, well-supported clade distinct from *Rickettsiella* species. They exhibit low ANI (84.29–88.19%) relative to their closest relative, *Diplorickettsia massiliensis* (shown in **a**), representing the first genomic record of this lineage in leafhoppers. GLMC_Bin048 branches separately, suggesting a divergent isolate within this novel group. **d**, Evolutionary placement of the deep-branching ‘*Ca.* Dalgrimnella’ lineage (GLMC_Bin066 and GLMC_Bin067). These highly divergent genomes lack close representatives in NCBI and occupy a phylogenetic position adjacent to the *Rickettsiales*, *Holosporales* and *Pelagibacterales*, representing a previously uncharacterized clade potentially influential in host biology.

### Plant-associated pathogenic and growth-promoting bacteria coexist within leafhopper microbiome assemblages

From the MAGs recovered, 88 were checked for potential beneficial or pathogenic plant interactions. These included taxa with representative species previously reported as potential plant pathogens, such as *Pseudomonas*, ‘*Candidatus* Phytoplasma’, *Pantoea*, *Spiroplasma*, *Enterobacter*, *Erwinia*, and other related genera. (**Figure 3d**, **Supplementary Table 10**). Consistent with this, genomes assigned to these lineages were enriched in genes associated with virulence, antibiotic resistance, and host interaction, and displayed high sequence identity to their reference pathogens (**Supplementary Figure 3**, **Supplementary Table 11**). These results indicate that leafhoppers harbour and potentially disseminate a broader spectrum of plant pathogens than previously recognized, extending beyond well-established agents such as phytoplasmas, *Spiroplasma*, and viruses. However, direct evidence of transmission will require targeted experimental validation.

Plant growth-promoting functions were also highly represented in leafhoppers’ microbiome, with 4,153 genes associated with nutrient acquisition, phytohormone modulation, and stress mitigation (**Supplementary Figure 3 and Figure 4**, **Supplementary Table 12)**. We further identified 143 orthologous genes associated with multiple bacterial secretion systems, including T1SS-T6SS, as well as the Sec, SRP, and Tat translocation systems **(Supplementary Table 13)**. Distinct secretion system profiles were observed among putative symbiont lineages, with enrichment of T3SS components in *Sodalis*, *Symbiopectobacterium*, and *Arsenophonus*, and predominant T4SS signatures in *Diplorickettsia*, *Rickettsia*, and *Wolbachia*. These patterns suggested divergent strategies of host interaction, with T3SS potentially mediating effector delivery and T4SS associated with intracellular persistence and host manipulation. The retention of diverse secretion systems may further reflect an evolutionary continuum from environmental or pathogenic bacteria toward symbiotic lifestyles, in which recently acquired or facultative symbionts may not yet have undergone extensive genome reduction or functional streamlining.

### Genome-resolved data reveal symbionts as multifunctional drivers of host ecology and evolution

Functional analyses on species-level MAG clusters (ANI > 95%) revealed distinct ecological and evolutionary stratification among obligate, secondary, and environmentally acquired symbionts (**Figure 5, Supplementary Table 14**). The leafhopper microbiome is predicted to contribute to host fitness by supplying essential and non-essential amino acids and vitamins, producing metabolites involved in abiotic stress alleviation and detoxification, and encoding functions associated with defense mechanisms, reproductive modulation, and the expansion of dietary capabilities through the degradation of novel food sources (**Figure 5**).

**Fig. 5.**
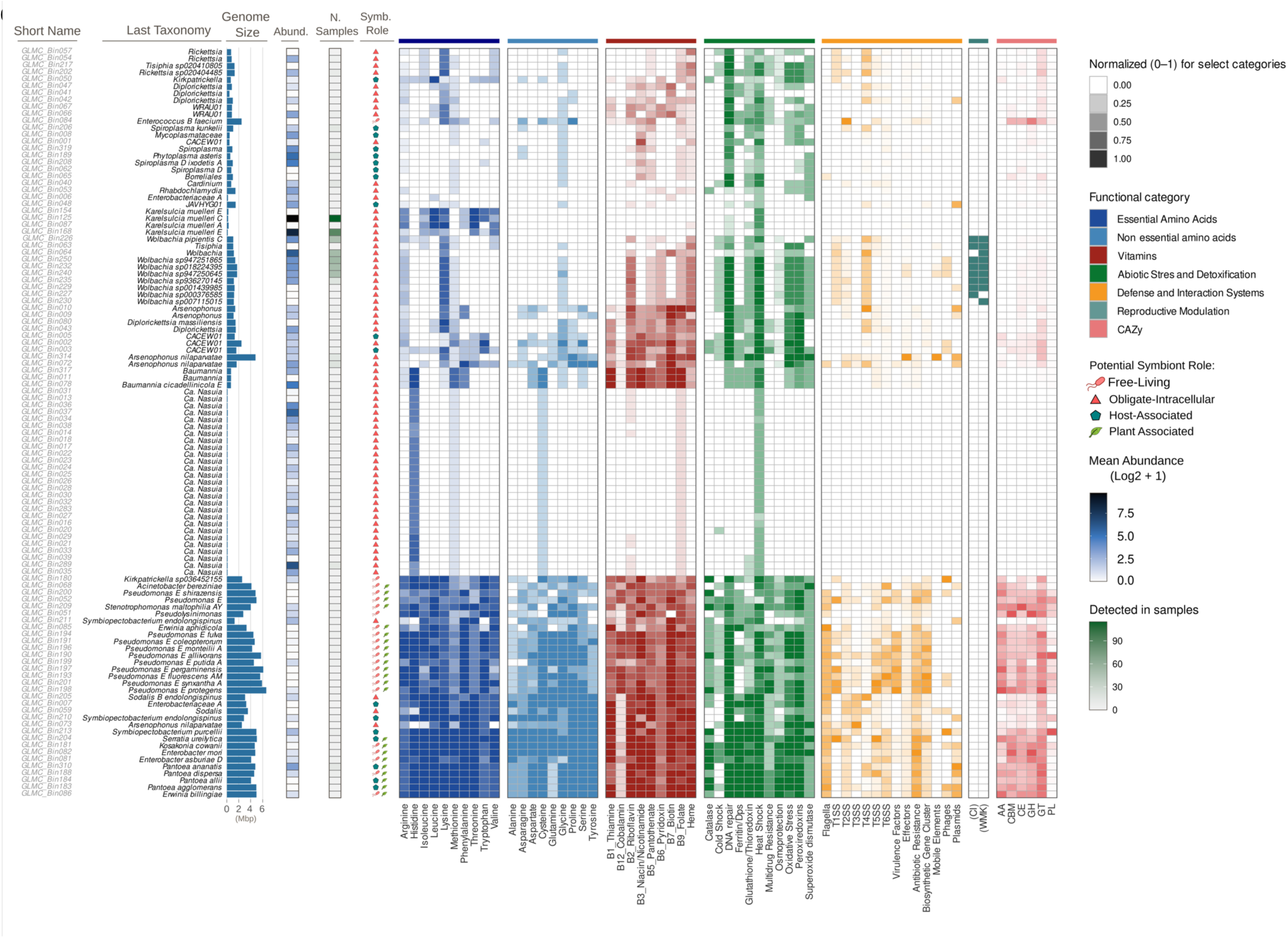
Genomic and functional landscape of the leafhopper microbiome. Heatmap displaying the metabolic potential of representative Metagenome-Assembled Genomes (MAGs) dereplicated at >95% Average Nucleotide Identity (ANI). **Rows** represent individual MAGs, annotated on the left by (from left to right): MAG identifier, lowest taxonomic assignment, genome size (Mbp), mean abundance across metagenomes (blue scale; Log2 + 1 transformed), sample prevalence (green scale; number of samples where the MAG was detected), and predicted symbiotic role. **Columns** represent functional categories, including amino acid and vitamin biosynthesis, abiotic stress response, defense/secretion systems, reproductive manipulation, and carbohydrate-active enzymes (CAZy). Tile intensity reflects the normalized presence or completeness (0–1) of genes/pathways within each functional category.

Obligate intracellular symbionts such as Karelsulcia and Nasuia exhibited highly streamlined functional repertoires dedicated to nutrient provisioning (**Figure 5, Supplementary Table 15-16**). Karelsulcia encoded biosynthetic pathways for six essential amino acids, complemented by Nasuia or ‘*Ca*. Baumannia’, which supplied histidine and methionine (**Figure 5)**. Consistent with its ancient and stable association with leafhoppers, Karelsulcia displayed a highly conserved genomic architecture (**Supplementary Figure 5**). Pangenome analysis, however, resolved host-associated microclusters across leafhopper species, forming two major clades that may correspond to Asian and American lineages, indicative of geographic structuring within Karelsulcia populations (**Supplementary Figure 5**). The Nasuia pangenome was highly variable, resolving into three distinct clusters and showing markedly lower genomic coherence, with a mean ANI of 82.85% compared with the higher genomic stability observed in Karelsulcia (average ANI = 96.86%) (**Supplementary Figure 5, Supplementary Table 17-18**). Nasuia genomic variability was most pronounced in *D. maidis*, where genomes were extremely reduced (n = 11), averaging 94.39 kb, representing some of the smallest Nasuia genomes reported to date. Consistent with advanced genome erosion, Nasuia genomes lacked key methionine biosynthesis genes (*metC* and *metE*), whereas Karelsulcia retained a relatively stable genome size of approximately 191 kb across hosts (**Supplementary Table 16**).

Additionally, other important and less characterized members of the leafhopper microbiome included *Rickettsia*, *Diplorickettsia*, *Wolbachia*, and WRA01. These taxa encoded biosynthetic pathways for amino acids such as arginine, isoleucine, lysine, methionine, and valine, as well as multiple B vitamins, cofactors, and functions associated with heat shock and oxidative stress resistance (**Figure 5**). All representative *Wolbachia* genomes recovered here encoded homologs of genes associated with cytoplasmic incompatibility (CI) and *Wolbachia* male killing (WMK), indicating a conserved genetic potential for reproductive manipulation. Another functional cluster was also formed by an underreported taxon, the newly identified ‘*Candidatus* Edelium’, and by well-known *Arsenophonus* and ‘*Ca*. Baumannia’, collectively encodes pathways for the biosynthesis of multiple amino acids, particularly arginine, lysine, methionine, and phenylalanine, and shows strong representation of B vitamin metabolism (**Figure 5**).

A final group identified in the functional heatmap corresponded to environmentally acquired bacteria, characterized by larger genome sizes (>2 Mb) and broad functional repertoires. This group included taxa such as *Serratia*, *Kosakonia*, *Enterobacter*, *Erwinia*, *Pantoea*, *Pseudomonas*, and *Pseudolysinimonas* (**Figure 5**). These organisms generally retained near-complete amino acid and vitamin biosynthetic capacities and encoded numerous genes related to abiotic stress tolerance. However, this functional breadth was accompanied by trade-offs, including expanded repertoires of antibiotic resistance genes, phage-associated elements, mobile genetic elements, and biosynthetic gene clusters, reflecting high genomic plasticity and frequent horizontal gene exchange. The presence of carbohydrate-active enzymes further suggests a potential role in degrading complex plant-derived compounds, possibly contributing to host nutrition or shaping gut microenvironments, as previously reported across a wide diversity of taxonomic groups and recently in the potato leafhopper, *Empoasca fabae* ^14–17^.

### Functional variation in symbiont assemblages of selected crop pests’ leafhoppers

To assess species- and geography-driven variation in microbiome structure and function, we analyzed two phloem-feeding leafhoppers with similar life strategies but distinct ecological ranges: *Macrosteles quadrilineatus* and *D. maidis*. Samples spanned a broad latitudinal gradient across the Americas, from Brazil to Nunavik, in the Canadian Arctic, allowing us to evaluate symbiont stability and environmental plasticity. In both species, the obligate symbionts Karelsulcia and Nasuia were detected in all samples (**Figure 6a to 6b**), forming a conserved metabolic core. Pangenome analyses revealed strong host species-specific structuring, with minimal influence of geographic distance, indicating that host identity is the primary driver of obligate symbiont genomic diversity **(Figure 6c)**.

**Fig. 6.**
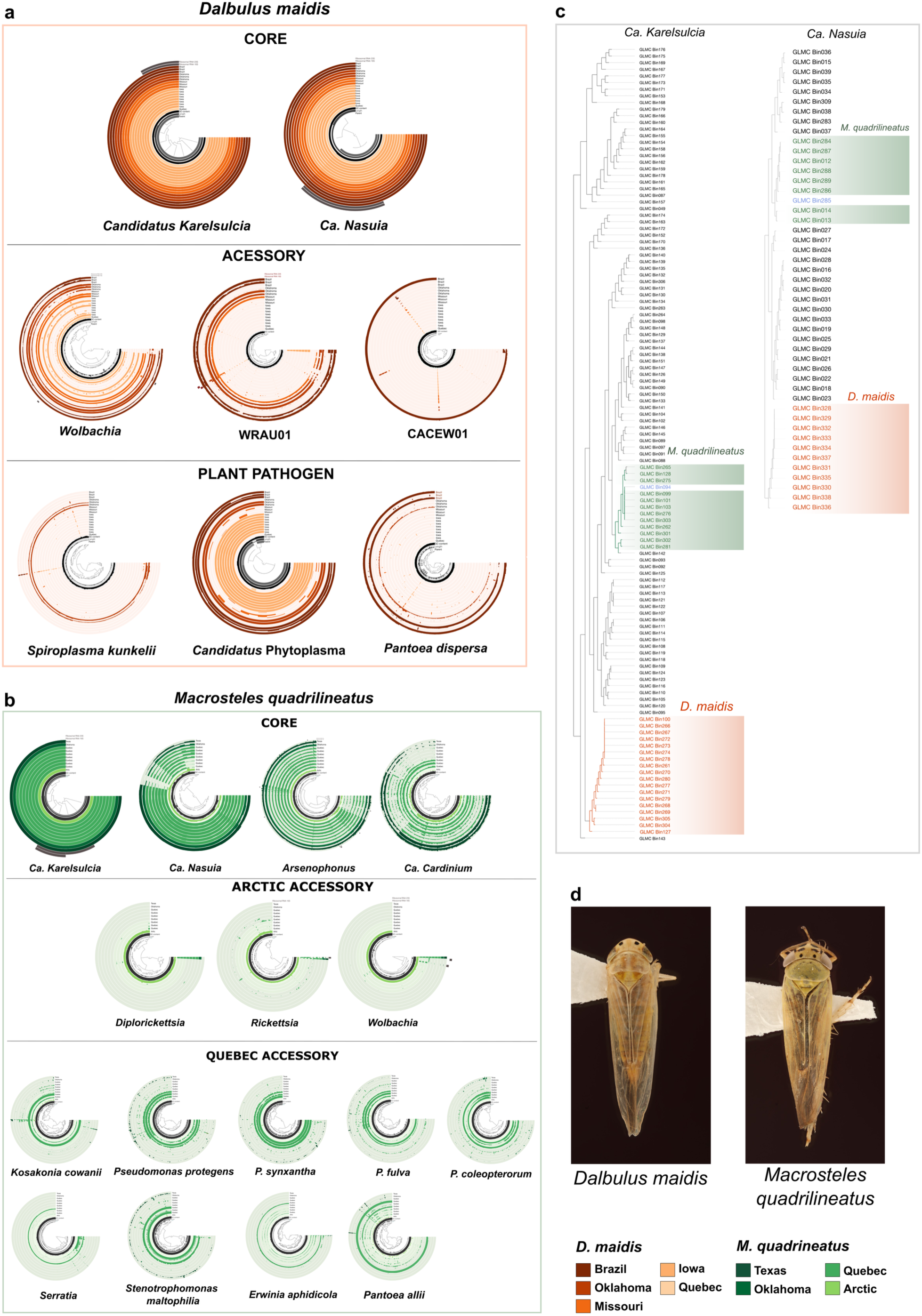
Biogeographical variability and core–accessory dynamics of the microbiome in key agricultural vectors. **a**, Comparative recruitment analysis for the corn leafhopper *Dalbulus maidis*, distinguishing the conserved core microbiome from accessory elements and putative plant pathogens. Variations in the coverage landscape highlight the differential acquisition of environmentally sourced bacteria across sampling sites.**b**, Metagenomic recruitment profiling of the *Macrosteles quadrilineatus* microbiome across distinct climatic zones (USA vs Arctic vs. Quebec). The Anvi’o display organizes microbial contigs (central dendrogram) based on differential coverage patterns. Concentric rings represent individual metagenomic samples; a continuous (filled) ring indicates the ubiquitous presence of a specific MAG (core microbiome), whereas discontinuous or fragmented rings highlight accessory taxa restricted to specific geographic subregions or low-abundance variants. **c**, Phylogenomic resolution of the primary obligate symbionts ‘*Ca*. Karelsulcia’ and ‘*Ca*. Nasuia’. The tree structure demonstrates strict host-species specificity, with symbiont lineages clustering congruently with host taxonomy (genus/subfamily) rather than geographic origin. Intraspecific branches show minimal divergence between distant populations, indicating that these co-obligate symbionts are evolutionarily conserved and do not provide sufficient resolution for geographic tracking of host populations. **d**, Representative images of the focal vector species: *Dalbulus maidis* (left) and *M. quadrilineatus* (right).

In *M. quadrilineatus*, *Arsenophonus* and *Cardinium* were consistently detected across North American populations, forming a stable symbiont core associated with nutritional provisioning and host regulation (**Figure 6b**). In contrast, *Diplorickettsia*, *Rickettsia*, and *Wolbachia* were detected exclusively in Canadian Arctic populations, suggesting context-dependent associations potentially linked to cold and environmentally harsh conditions (**Figure 6b**). This pattern is supported by the enrichment of cold-adaptation-related genes, including cold shock proteins, RNA helicases, lipid-modification pathways, and systems involved in compatible solute transport and redox homeostasis (**Supplementary Figure S7, Supplementary Table S19)**. Notably, the Arctic *Arsenophonus* MAG encoded an expanded repertoire of cold-tolerance-associated genes (**Supplementary Figure 7**). All *Wolbachia* MAGs recovered from Arctic *M. quadrilineatus* and *Macustus grisescens,* another leafhopper found in the arctic^18^ encoded homologs of the CI and WMK genes, indicating a conserved genetic potential for reproductive manipulation even in high-latitude populations. In addition, Québec populations harbored a broader diversity of environmentally acquired taxa, including *Kosakonia*, *Pseudomonas*, *Stenotrophomonas*, *Erwinia*, and *Pantoea* (**Figure 6b**).

The corn leafhopper, *D. maidis,* displayed a simpler but more variable symbiotic repertoire than *M. quadrilineatus* (**Figure 6c**). While Karelsulcia and Nasuia also formed a conserved core, *Wolbachia* showed an uneven distribution across samples from Brazil, the United States, and Canada. All *Wolbachia* MAGs from *D. maidis* contained homologs of CI genes and high-identity members of the WMK gene family. A *Pantoea dispersa* MAG was detected only in Brazilian samples, indicating region-specific acquisition, whereas the undescribed lineage WRA01 was consistently recovered across distant regions, supporting its status as a stable, emerging symbiont (**Figure 6a**). Plant pathogens *Spiroplasma kunkelii* and ‘*Ca.* Phytoplasma’ were also detected, highlighting the coexistence of obligate, facultative, and pathogenic microbes within *D. maidis* populations.

## DISCUSSION

In this study, we introduce the Global Leafhopper Microbiome Catalog (GLMC) as a genome-resolved resource to investigate symbiosis, ecological interactions, and microbial evolution in leafhoppers, one of the most diverse and agriculturally important insect groups^19^. Across 171 species spanning 11 subfamilies and 13 countries, genome-resolved metagenomics revealed a recurrent, tripartite stratification of microbial associates that largely transcends host phylogeny and geographic origin. This organization is consistent with a modular microbiome architecture in which symbiont communities are structured by evolutionary depth, transmission mode, and degree of functional integration with the host ^4,20–23^.

At the core of this architecture lie obligate and co-obligate symbionts that exhibit long-term co-evolution and deep metabolic interdependence with their hosts. These bacteria, principally Karelsulcia and Nasuia, supply essential amino acids that are indispensable for leafhopper development and survival on nutrient-limited phloem sap^2,4,24^. Their genomic repertoires are highly streamlined and conserved across host lineages, consistent with ancient, vertically transmitted associations^2,4,7,24^. Pangenome analyses, however, revealed host-associated geographic structuring within Karelsulcia, distinguishing American from Asian lineages, while Nasuia displayed markedly greater genomic variability, including extremely reduced genomes in *D. maidis* that had lost key methionine biosynthesis genes (*metC* and *metE*). These patterns suggest that even the most deeply integrated symbioses retain detectable evolutionary plasticity shaped by host ecology and lineage-specific selective pressures.

An intermediate layer of facultative symbionts, including *Wolbachia*, *Arsenophonus*, *Rickettsia*, *Cardinium*, *Diplorickettsia*, and *Tisiphia*, exhibited heterogeneous distributions across host lineages, with distinct leafhopper subfamilies harboring partially overlapping but non-identical symbiont assemblages. Closely related symbiont lineages were recovered in distantly related host clades, while sister host species often differed in their secondary symbiont complements. This pattern argues against strict host-symbiont co-diversification and instead supports a model of recurrent, temporally independent acquisition followed by differential retention, as recently reported for *M. quadrilineatus*^25^. The repeated recruitment of functionally comparable secondary symbionts across independent leafhopper lineages suggests that these bacteria occupy conserved adaptive niches within the host^7,11,26^. Their intermediate genome sizes, retention of secretion systems, and metabolic flexibility are hallmarks of symbionts that likely modulate host physiology, environmental stress tolerance, and ecological interactions rather than providing strictly essential nutrients, consistent with the emerging view of insect symbioses as a dynamic continuum in which facultative associates shift between mutualism, commensalism, and parasitism depending on ecological context^20,27–30^. Notably, *Wolbachia* genomes recovered here encoded homologs of cytoplasmic incompatibility and male-killing loci, indicating a conserved genetic potential for reproductive manipulation across host species and geographic ranges, including high-latitude Arctic populations. A similar layered organization, structured around obligate nutritional symbionts, recurrent secondary symbionts, and environmentally acquired associates, has been described in dipterans and other hemipterans, reinforcing the view that this architecture reflects transmission mode, genome reduction, and functional specialization rather than host phylogeny alone^22,31–35^.

The outermost layer of the leafhopper microbiome is dominated by environmentally acquired bacteria that exhibit rapid taxonomic turnover and weak phylogenetic constraint, reflecting frequent horizontal acquisition from plants, soil, and water. These taxa, including *Pseudomonas*, *Erwinia*, *Pantoea*, *Enterobacter*, *Kosakonia*, and *Serratia,* retain broad functional repertoires encompassing amino acid and vitamin biosynthesis, abiotic stress tolerance, and carbohydrate-active enzyme activity, but their association with the host is facultative and does not constitute an essential symbiotic dependency. Similar environmentally sourced microbial layers have been described in other insects and are thought to enhance functional breadth at the community level while remaining evolutionarily decoupled from the host^36–39^. The recurrent detection of known vector-borne pathogenic bacteria, including ‘*Ca.* Phytoplasma’, and, *S. kunkelii*, together with other bacteria with plant-associated lineages such as *Pantoea* and *Erwinia* within leafhopper microbiomes reinforce the role of these insects as ecological connectors among microbial reservoirs in plants, soil, and insect vector populations. These findings suggest that, beyond well-established transmission of phytoplasmas and spiroplasmas, leafhoppers may harbour and potentially facilitate the movement of a broader spectrum of plant-associated microbes than currently recognized, a hypothesis that warrants evaluation through targeted transmission assays^40–43^.

Beyond the known microbial diversity, the GLMC resolved several novel bacterial lineages lacking close relatives in existing genomic databases. These include a putative new family within Thiotrichales (CACEW01), consistently recovered across multiple host species, including the major crop pests *D. maidis* and *Graminella nigrifrons*; a Diplorickettsia-like clade (ANI 84–88% relative to *D. massiliensis*) representing the first genomic characterization of this group in leafhoppers; and two deep-branching genomes (WRA01) phylogenetically adjacent to Rickettsiales, consistently recovered across geographically distant host populations, supporting their status as stable, emergingsymbionts. These discoveries underscore the extent to which leafhopper-associated microbiomes remain genomically uncharted and highlight the value of large-scale, culture-independent approaches for uncovering novel microbial diversity.

Genome-resolved approaches such as the GLMC enable precise discrimination between persistent symbionts and transient taxa and allow bacterial lineages to be traced along the mutualism-to-pathogenicity continuum. This resolution is particularly critical in agricultural systems, where leafhoppers are major vectors of plant pathogens, and microbiome composition can modulate vector competence and disease emergence. Moreover, by expanding genomic representation within an insect family whose microbiota remains poorly captured in current databases, the GLMC helps address a major gap in genome-resolved resources for host-associated microbial diversity. By providing a comprehensive genomic framework for leafhopper-associated microorganisms, the GLMC establishes a foundation for investigating symbiont-mediated host adaptation, microbiome-informed pest management, and the ecological and evolutionary processes that shape host–microbe interactions.

## METHODS

### Sample collection and metagenomic data sources

In this study, data were sourced from two distinct origins: (*i*) 38 original metagenomic data generated from field-collected insect samples across multiple regions and countries; and (*ii*) 214 publicly available metagenomic datasets retrieved from NCBI. The field-collected samples primarily included *Dalbulus maidis*, *Empoasca fabae*, and *M. quadrilineatus*; and *Diplocolenus abdominalis*, *Hebecephalus algidus*, *Limotettix myralis*, *Limotettix plutonious*, *Macustus grisescens*, *Psammotettix allienus*, and *Strogylocephalus mixtus* from the Canadian Arctic (**Supplementary Table S1**). These collections represent diverse geographic locations, including Canada, the United States, and Brazil. Insects were sampled using entomological aspirators and yellow sticky traps, following the protocol previously established^1^. To reduce surface contamination, specimens were washed with sterile phosphate-buffered saline (PBS) containing 0.01% Tween-20. Genomic DNA was extracted as previously described^44^. DNA library preparation, the Nextera XT DNA Library Preparation Kit (Illumina, USA) was used according to the manufacturer’s instructions. Sequencing was performed on the Illumina NovaSeq 6000 platform with paired-end 150 bp reads, yielding high-resolution data suitable for comprehensive taxonomic and functional characterization of insect-associated microbiomes.

### Publicly available metagenomes download and curation

Publicly available datasets used here were primarily generated for purposes unrelated to microbiome research, such as the assembly of mitochondrial or insect genomes (**Supplementary Table S1**). Here, we repurposed these datasets for microbiome analysis, recognizing their potential to provide insights into insect-associated microbial communities. Data were retrieved from the Sequence Read Archive (SRA) hosted by NCBI. We systematically searched for metagenomic datasets derived from whole leafhopper (Hemiptera: Cicadellidae) specimens, focusing specifically on untargeted (shotgun) sequencing approaches. To ensure an unbiased representation of microbial communities, we excluded studies that employed targeted sequencing strategies, such as amplicon-based 16S rRNA profiling, anchored hybrid enrichment, or other locus-specific methods. Both long-read and short-read sequencing datasets were included: long-read platforms such as Oxford Nanopore Technologies and Pacific Biosciences (PacBio), and short-read technologies like Illumina HiSeq, MiSeq, and NovaSeq. All available raw data files were downloaded using the prefetch and fasterq-dump utilities from the SRA Toolkit (v3.0.0) to ensure data integrity and compatibility with downstream processing.

### Quality control, host-removal, and metagenomic assembly

Original and public available data were then evaluated for quality control with fastqc and were quality-trimmed using fastp v.0.23.4^45^ to keep only high- to moderate quality reads (HQ) (Q > 20). A first taxonomic classification was performed with Kraken2 v.2.1.3 against the core-nt database^46^ and with SingleM against the GTDB database^47^. To eliminate host-derived reads (e.g., from leafhoppers, humans, and plants), we used KrakenTools v.1.2 (https://github.com/jenniferlu717/KrakenTools). Filtered reads were then assembled using two metagenomic assemblers, MEGAHIT v.1.1.4^48^ and MetaSPAdes v.4.1.0^49^ for short reads, to compare performance and maximize contig recovery. Long-read sequences were assembled using MetaFlye^50^.

### Gene prediction, classification, and clustering

To facilitate sample tracking, contig files from each sample were initially renamed to include sample-specific identifiers. All assembly files were concatenated into a single *fasta* file that contained all contigs from all samples. From this file ribosomal RNA genes were extracted using Barrnap v.0.9. Then, rRNAs were taxonomically classified with blastn with previously downloaded genbank and refseq genomes (on 15 august 2025). These sequences were used to delimitate possible species present in leafhopper microbiome, and to be used as target species in metagenome-assembled genome retrieval but also for built phylogenetic trees. For gene catalog construction, open reading frames (ORFs) were predicted using Prodigal (v2.6.3)^51^ with the -p meta parameter, optimized for metagenomic data. ORFs shorter than 100 base pairs were filtered out to avoid potential artifacts. The remaining ORFs were clustered using MMseqs2 (v15.6f452)^52^ with the parameters --min-seq-id 0.95 and -c 0.8, resulting in a nonredundant microbial gene catalog by collapsing highly similar sequences.

### Taxonomic and functional annotation of ORFs and downstream analyses

ORFs from the gene catalog were re-classified using Kraken2 with the Core-nt database. Functional annotation was performed using eggNOG-mapper v2.1.2^53^ against the eggNOG database version 5.0.2^54^. The eggNOG database integrates functional annotations from several databases, including KEGG, the Carbohydrate-Active Enzymes (CAZy) database, and the Cluster of Orthologous Groups (COG) categories.

For quantifying gene abundance among different leafhopper species, reads from each sample were aligned to the gene catalog using Salmon v.1.10^55^. Gene abundance profiles were calculated in transcripts per million (TPM), accounting for both gene length and total mapped reads per sample. Taxonomic, functional, and coverage data were integrated and processed in the microeco R package for downstream analyses^56^. Prior to taxonomic profiling, data were rarefied to the lowest sequencing depth across samples to ensure comparability. Analyses were conducted at both the gene level (individual ORFs per treatment) and genus level (aggregated taxonomy). Microbial community structure was evaluated using Principal Coordinate Analysis (PCoA) based on Bray-Curtis dissimilarity, with statistical significance assessed via PERMANOVA (p < 0.05).

### Metagenome binning and genome quality control

To enable robust and accurate taxonomic and functional analyses, we employed a metagenomic binning strategy to reconstruct metagenome-assembled genomes (MAGs). All steps were implemented using MAG_Tools, a software package with dedicated modules for recovering, evaluating, and cleaning MAGs developed for this study (***Github: Edelab***). To maximize genome recovery, we applied four complementary binning algorithms in MAG_Tools –w MAG_finder: MetaBAT2 (v2.12.1)^57^, MaxBin2^58^ (v2.2.7), CONCOCT^59^, and VAMB^60^. In addition, MAG_Tools integrates a 16S rRNA-guided binning procedure (MAG_rRNA), in which rRNA genes predicted by Barrnap served as taxonomic anchors. This was useful in some MAGs that was not recovered by using classic binning tools (i.e. ‘*Candidatus Nasuia’*). For this, we conducted the genome-guided binning by aligning contigs to reference genomes with BLASTn, followed by contig recruitment using SeqKit^61^. This manual refinement allowed the recovery of several symbionts that were not detected by automated binning approaches. Further refinement of MAGs was performed using DAS_Too^62^, and redundancy was reduced via dereplication with dRe^63^. MAG quality was assessed using a multidimensional framework encompassing structural, functional, and taxonomic evaluations: MAG_sumary. Structural features were analyzed with QUAST^64^, functional annotation was performed with Prokka^65^, and taxonomic composition of contigs was evaluated with GUNC^66^. Genome completeness and contamination were estimated using both CheckM^67^ and CheckM2^68^. MAGs were manually curated to keep non-redundant representatives for each leafhopper species. MAGs with higher than 1,000 contigs were removed, and N50 and completeness were used to estimate the best MAGs for each species. We then removed MAGs that have low number of tRNAs (tRNAs < 10) and Coding Sequences (CDS < 100).

### Taxonomic annotation, phylogenetic tree construction, functional annotation, and abundance analysis

All MAGs were taxonomically classified using GTDB-Tk (v2.3.2)^69^ with the Genome Taxonomy Database (GTDB) release 226^70^, providing standardized and high-resolution taxonomic assignments. To further validate taxonomic identities and reduce classification bias, we also applied GUNC, offering an independent assessment of MAG taxonomy and contamination. We considered a species to represent a novel taxon when it lacked a representative in the GTDB, as defined by ANI below 95.

Phylogenomic relationships among MAGs were inferred using GToTree^71^ with the -H Bacteria parameter to select bacterial marker genes, and the resulting tree was visualized in iTOL. We performed an additional step to verify the taxonomic composition and internal consistency of MAG contigs using Anvi’o^72^, examining contig-level coverage across samples. This approach helped discriminate genuine symbionts or host-associated bacteria from possible chimeric assemblies in in which contigs display highly divergent taxonomies. For MAGs detected in a single sample, we inspected the assemblies in anvi-interactive to determine whether contigs, exhibited coherent, uniform coverage patterns, as expected for authentic genomes, or heterogeneous patterns suggestive of contamination.

Functional annotation of MAGs was conducted with Prokka, enabling comprehensive gene annotation and exploration of pathways related to diverse metabolic functions, stress responses, and ecological interactions. We also used anvi-estimate-metabolism to evaluate the completeness of the metabolism in each MAG. CoverM^73^ was used to quantify the relative abundance and sequencing depth of each MAG across all metagenomic samples by mapping quality-filtered reads back to the MAG assemblies. Based on these considerations, we generated a presence–absence matrix by defining MAG presence when normalized abundance, measured as Reads Per Kilobase per million mapped reads (RPKM), was equal to or greater than 1.

### Inference of host interaction, virulence, and metabolic adaptation

To identify genes associated with pathogenicity and plant interactions, we performed a detailed functional annotation. We first annotated the MAGs using *PGPg_finder* with the PLaBaSe database to detect genes potentially related to virulence or plant-associated traits^74–76^. These results were validated by screening the same targets within the Prokka and Anvi’o annotations. The identified genes were then categorized into functional groups, including motility, secretion systems, and related traits. Virulence factors were annotated using the Abricate tool (https://github.com/tseemann/abricate) with the VFDB database^77^, and effector or pathogen-host interaction genes were identified through the PHI-Base database^78^. Antibiotic resistance genes were characterized using the CARD database^79^. Plasmids were predicted using PlasmidFinder^80^, and mobile genetic elements were confirmed by searching for integrases, resolvases, recombinases, and transposases in the annotation outputs. Genes associated with host reproductive modulation were screened, including loci involved in *Wolbachia-*mediated cytoplasmic incompatibility and male killing. Reference sequences for these targets were downloaded from NCBI and queried against the MAGs using BLAST. To investigate secondary metabolism, we performed biosynthetic gene cluster analysis using antiSMASH version 6.1.1^81^ with default parameters, enabling the detection of established and putative clusters involved in natural product biosynthesis. Carbohydrate-active enzymes were annotated in Anvi’o using the CAZy database^82^.

## DATA AVAILABILITY

All datasets of metagenomic samples used in this study are available in NCBI under the codes provided in Sample S1. The gene catalog under 95% are available at Zenodo https://zenodo.org/records/20042173 with gene annotation and taxonomy available for each similarity level. MAGs provided in this study were deposited in NCBI under BioProject PRJNA1450707 and are also available in Zenodo https://zenodo.org/records/20042173. All MAG annotations are available in Zenodo. Intermediate analyses and files are available as compressed archives on Zenodo.

## CODE AVAILABILITY

Code sources of analyses used in this study are available on GitHub EdeLab (https://github.com/Edelab/MAG_Tools). The MAG_Tools, which is a pipeline used exclusively in this study, are also provided in Github.

## ACKNOWLEDGEMENTS

This work was funded by the Réseau québécois de recherche en agriculture durable (RQRAD), the Ministère de l’Agriculture, des Pêcheries et de l’Alimentation du Québec (MAPAQ), and the Fonds de recherche du Québec–Nature et technologies (FRQNT) through the Programme de recherche en partenariat—Agriculture durable—Volet II—2e concours (application number 337847), as well as by the Natural Sciences and Engineering Research Council of Canada (NSERC) through the Alliance-SARI Program (Grant ALLRP 588519-23). MCC thanks for the Fundação de Amparo à Pesquisa e Inovação do Estado de Santa Catarina-Brazil [grant no FAPESC2023TR000218]. The Nunavik Sentinels program from the Montreal Insectarium thanks all Nunavik and Eeyou Istchee collaborators and participants. This program was funded in part by RBC through Espace pour la vie Foundation.

## AUTHOR CONTRIBUTIONS

E.P.-L. designed and supervised the research. N.P., J.J., A.G.-T., M.C.C., M.R.D., A.M.F., A.O.V., and I.V. collected samples. J.M. prepared materials for sequencing. T.A.P. performed genome assembly and annotation and performed microbiome analysis and interpretation. T.A.P., A.A.S., J.M., and E.P.-L. wrote the manuscript. T.A.P. prepared the figures and tables. T.A.P., J.M., A.A.S., N.P., J.J., A.G.-T., M.C.C., M.R.D., A.M.F., A.O.V., I.V., and E.P.-L. edited the article.

## COMPETING INTERESTS

The authors declare no competing interests.

## ADDITIONAL INFORMATION

**Fig S1 | Distribution of host leafhopper taxa across the Global Leafhopper Microbiome Catalog.** Horizontal bar chart showing the frequency of all host genera represented in the 264 sequenced leafhopper datasets. This expanded view complements Fig. 1 by resolving rare genera and highlighting the highly uneven contribution of host lineages, with a subset of genera accounting for most samples in the dataset.

**Fig. S2 | Host phylogenetic signal in bacterial community structure based on 16S rRNA.** Principal Coordinate Analysis (PCoA) of 16S rRNA amplicon profiles from 264 leafhopper microbiomes colored by host subfamily. PERMANOVA statistics indicate that bacterial community composition exhibits increasing explanatory power at finer host taxonomic levels, with the strongest association observed at the species level. This supports the hypothesis that host evolutionary history partially constrains microbiome structure.

**Fig. S3 | Functional profiling and hierarchical clustering of potentially pathogenic MAGs.** The dendrogram (left) groups Metagenome-Assembled Genomes (MAGs) based on PGP profiles, selecting for genera with known pathogenic members or suspected virulence potential. The heatmap displays annotation results across five categories: VFDB (Virulence Factor Database), CARD (Comprehensive Antibiotic Resistance Database), PHI-base (Pathogen-Host Interaction database), PGP (PLaBAse), and Secretion Systems (Type I–VI). For the first four categories, paired columns indicate the total number of gene hits (left sub-column) and the mean percentage identity (right sub-column). The Secretion Systems panel shows the count of identified components. Tile color intensity correlates with the magnitude of the values, which are explicitly annotated within each cell for clarity.

**Fig. S4 | Presence-absence profile of secretion system genes.** A binary heatmap displaying the specific gene repertoire for bacterial Secretion Systems I through VI across representative Metagenome-Assembled Genomes (MAGs) in the Global Leafhopper Microbiome Catalog (GLMC). The Y-axis lists the MAGs (arranged by hierarchical clustering), while the X-axis corresponds to individual genes encoding structural and effector components. Tile color indicates genomic conservation: red represents the presence of the specific gene, and white denotes its absence.

**Fig. S5 | Comparative pangenomics of the obligate symbionts ‘*Candidatus* Karelsulcia’ and ‘*Ca*. Nasuia’.** Pangenome analysis of the two dominant obligate symbionts, (Left) ‘*Ca*. Karelsulcia’ and (Right) ‘*Ca*. Nasuia’. In these circular plots, radial layers (rings) represent individual MAGs, and radial rays represent gene clusters. Dark/black segments indicate the presence of a gene cluster, while light/gap segments indicate absence. Genomes are organized by hierarchical clustering based on gene cluster distribution (central dendrogram). Inner metadata rings distinguish the host source: MAGs derived from *Macrosteles* species are colored green, while those from *Dalbulus* are orange. The inset heatmap (top right of each circle) displays pairwise Average Nucleotide Identity (ANI) values, highlighting genomic similarity alongside gene content. Outer layers summarize gene cluster statistics, including single-copy core genes (SCG), geometric and functional homogeneity, and maximum number of paralogs. Green distinct rings in the outer layers mark clusters with functional annotations in Pfam, KEGG, or COG databases. The comparison reveals that the ‘*Ca*. Karelsulcia’ pangenome is significantly more stable and conserved than the structurally variable ‘*Ca*. Nasuia’ pangenome.

**Fig. S6 | Distribution of abiotic stress tolerance genes in Arctic leafhopper microbiomes.** Binary heatmap profiling the presence of genes conferring resistance to environmental stressors in Metagenome-Assembled Genomes (MAGs) recovered from the Canadian Arctic (Nunavik). Rows correspond to specific stress response genes, while columns represent individual MAGs. Purple tiles indicate the presence of a gene, and white tiles denote its absence. Green circles highlight MAGs derived from *Macrosteles* host populations as well as the *Arsenophonus* lineage, distinguishing them from other recovered taxa.

**Fig. S7 | Prokaryotic taxonomic profiling across leafhopper lineages.** A binary heatmap displaying the distribution of bacterial genera identified via SingleM profiling using the GTDB marker gene database. Rows represent bacterial taxonomy (clustered hierarchically), while columns represent leafhopper host genera, grouped by their respective subfamilies (indicated by the top color bar). Red tiles indicate the presence of a bacterial genus, and white tiles denote its absence. The dense cluster at the bottom highlights the widespread prevalence of obligate symbionts, including ‘*Candidatus* Karelsulcia’, ‘*Ca*. Nasuia’, and ‘*Ca*. Baumannia’, across diverse host lineages.

**Fig. S8 | Prokaryotic taxonomic profiling across leafhopper lineages.** Heatmap displaying the standardized relative abundance (Z-scores) of bacterial genera identified via SingleM profiling (GTDB). Rows represent hierarchically clustered bacterial taxa, while columns represent leafhopper host genera grouped by subfamily (indicated by the top color bar). Tile color intensity reflects the Z-score, where deeper red indicates higher relative abundance compared to the row mean. The analysis reveals a distinct high-abundance cluster at the bottom comprising obligate symbionts ‘*Candidatus* Karelsulcia’, ‘*Ca.* Nasuia’, and ‘*Ca.* Baumannia’, which contrasts with the upper or more variable abundance of facultative and environmental taxa in the lower clusters.

**Table S1 | Metadata and sequencing information for leafhopper metagenomes.** Detailed list of the leafhopper samples analyzed in this study, including host taxonomy (Subfamily, Tribe, Species), geographic origin, sequencing platform specifications, and corresponding NCBI BioProject/SRA accession numbers.

**Table S2 | Metrics of the non-redundant microbial gene catalog.** Assembly statistics and global properties of the constructed gene catalog, detailing the total number of non-redundant DNA sequences, Open Reading Frames (ORFs), average sequence lengths, and dereplication results.

**Table S3 | Taxonomic classification of the gene catalog.** Taxonomic profiling of the microbial ORFs based on Kraken2 classification against the NCBI database. The table presents the distribution of reads and percentages across major taxonomic ranks (Kingdom to Species).

**Table S4 | Taxonomic marker gene abundance profiles.** Quantification of specific taxonomic marker genes (including 16S rRNA gene fragments) recovered from the metagenomes. Abundances are normalized as Transcripts Per Million (TPM) and classified by Phylum and Genus.

**Table S5 | Global functional annotation statistics.** Summary of the functional annotation of the gene catalog using EggNOG-mapper. Data includes gene counts and percentages assigned to major functional databases: COG categories, KEGG (Pathways, Modules, KO), CAZy (Carbohydrate-Active Enzymes), and Pfam domains.

**Table S6 | Metabolic landscape of the microbiome (KEGG Level 2).** Cumulative abundance (TPM) of genes categorized into KEGG Level 2 metabolic pathways. Highlights key functional guilds such as “Amino acid metabolism,” “Metabolism of cofactors and vitamins,” and “Xenobiotics biodegradation.“

**Table S7 | Comprehensive catalog of recovered MAGs and quality metrics.** Master table providing detailed metrics for all Metagenome-Assembled Genomes (MAGs). Includes taxonomic assignment (GTDB-Tk), genome quality assessment (CheckM completeness/contamination, N50, size), host source, and representative status.

**Table S8 | Distribution of MAGs across host lineages.** Summary statistics showing the count of recovered MAGs stratified by leafhopper host subfamily and by bacterial genus. This table highlights the prevalence of major symbiont lineages (e.g., *Sulcia, Nasuia*) across the sampled biodiversity.

**Table S9 | Taxonomic novelty and ANI divergence.** Analysis of Average Nucleotide Identity (ANI) revealing potential taxonomic novelty. Lists MAGs that likely represent undescribed species, genera, or families based on genomic divergence from reference databases.

**Table S10 | Lifestyle and pathogenicity classification.** Categorization of representative MAGs based on inferred lifestyle and pathogenic potential. Genomes are labeled as insect endosymbionts, potential plant pathogens (e.g., *Candidatus* Phytoplasma), or opportunistic pathogens based on genomic signals.

**Table S11 | Virulence factors and resistome profiling.** Genomic screening for pathogenic traits and antimicrobial resistance. The table lists gene hits against the VFDB (Virulence Factor Database), PHI-base (Pathogen-Host Interaction), and CARD (Antibiotic Resistance) databases, including mean identity percentages.

**Table S12 | Plant Growth-Promoting (PGP) traits inventory.** Presence-absence profile of genes associated with plant beneficial functions. Includes genes involved in nutrient solubilization, phytohormone modulation, and biocontrol mechanisms across the recovered MAGs.

**Table S13 | Bacterial secretion systems and competitive exclusion.** Detailed annotation of secretion system components (Types I through VI) and genes involved in competitive exclusion (e.g., conjugation machinery). Lists specific gene identifiers (e.g., *traI, comB9*) detected in each MAG.

**Table S14 | Functional genomic profile across GLMC MAGs.** Comprehensive inventory of metabolic pathways, stress response systems, interaction mechanisms, and mobile genetic elements identified across all non-redundant MAGs in the Global Leafhopper Microbiome Catalog (GLMC). Rows represent functional categories, including amino acid and vitamin biosynthesis, stress tolerance mechanisms, secretion systems, virulence-associated traits, and carbohydrate-active enzymes, while columns correspond to individual MAGs. Values indicate gene counts associated with each function per genome, providing a genome-resolved view of functional potential and enabling comparative assessment of metabolic capabilities, host interaction strategies, and adaptive traits across leafhopper-associated microbiomes.

**Table S15 | Pangenome of Sulcia-associated genomes.** Pangenomic reconstruction generated with Anvi’o integrating all GLMC MAGs assigned to ‘*Candidatus* Sulcia’ and publicly available reference genomes. The table summarizes gene cluster distribution across genomes, highlighting core, accessory, and unique gene sets, and enabling comparative assessment of genomic conservation, functional partitioning, and lineage-specific adaptations within this obligate nutritional symbiont.

**Table S16 | Pangenome of Nasuia-associated genomes.** Pangenomic reconstruction generated with Anvi’o integrating all GLMC MAGs assigned to “*Candidatus* Nasuia” and publicly available reference genomes. The table details gene cluster presence across genomes, allowing discrimination of conserved core functions and variable accessory repertoires, and providing insights into genome reduction patterns, functional specialization, and evolutionary divergence within this highly streamlined symbiont lineage.

**Table S17 | Pairwise ANI values among Sulcia-associated genomes.** Matrix of average nucleotide identity (ANI) calculated between all MAGs assigned to “*Candidatus* Sulcia” and reference genomes. Values represent genome-wide nucleotide similarity between pairs of genomes, enabling assessment of genomic relatedness, species boundaries, and population structure within this conserved symbiont lineage.

**Table S18 | Pairwise ANI values among Nasuia-associated genomes.** Matrix of average nucleotide identity (ANI) calculated between all MAGs assigned to “*Candidatus* Nasuia” and reference genomes. Values indicate genome-wide similarity across pairs, providing resolution of genomic divergence, strain-level variation, and evolutionary relationships within this highly reduced symbiont clade.

**Table S19 | Abiotic stress tolerance profile.** Genomic inventory of genes conferring resistance to environmental stressors. The table indicates the presence or absence of molecular chaperones (e.g., *dnaK, groL*) and membrane modification genes associated with thermal and osmotic stress response.

## Notes

### Competing Interest Statement

The authors have declared no competing interest.

