## Supplementary Figures for "Genome-resolved metagenomics reveals conserved, flexible and emerging symbioses across global leafhoppers"

### ADDITIONAL INFORMATION

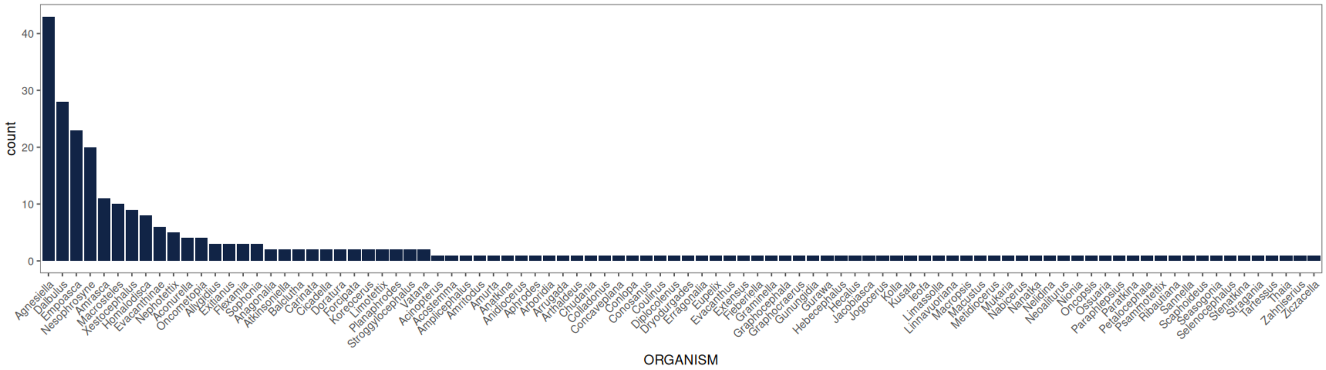


**Fig S1 | Distribution of host leafhopper taxa across the Global Leafhopper Microbiome Catalog.** Horizontal bar chart showing the frequency of all host genera represented in the 264 sequenced leafhopper datasets. This expanded view complements Fig. 1 by resolving rare genera and highlighting the highly uneven contribution of host lineages, with a subset of genera accounting for most samples in the dataset.

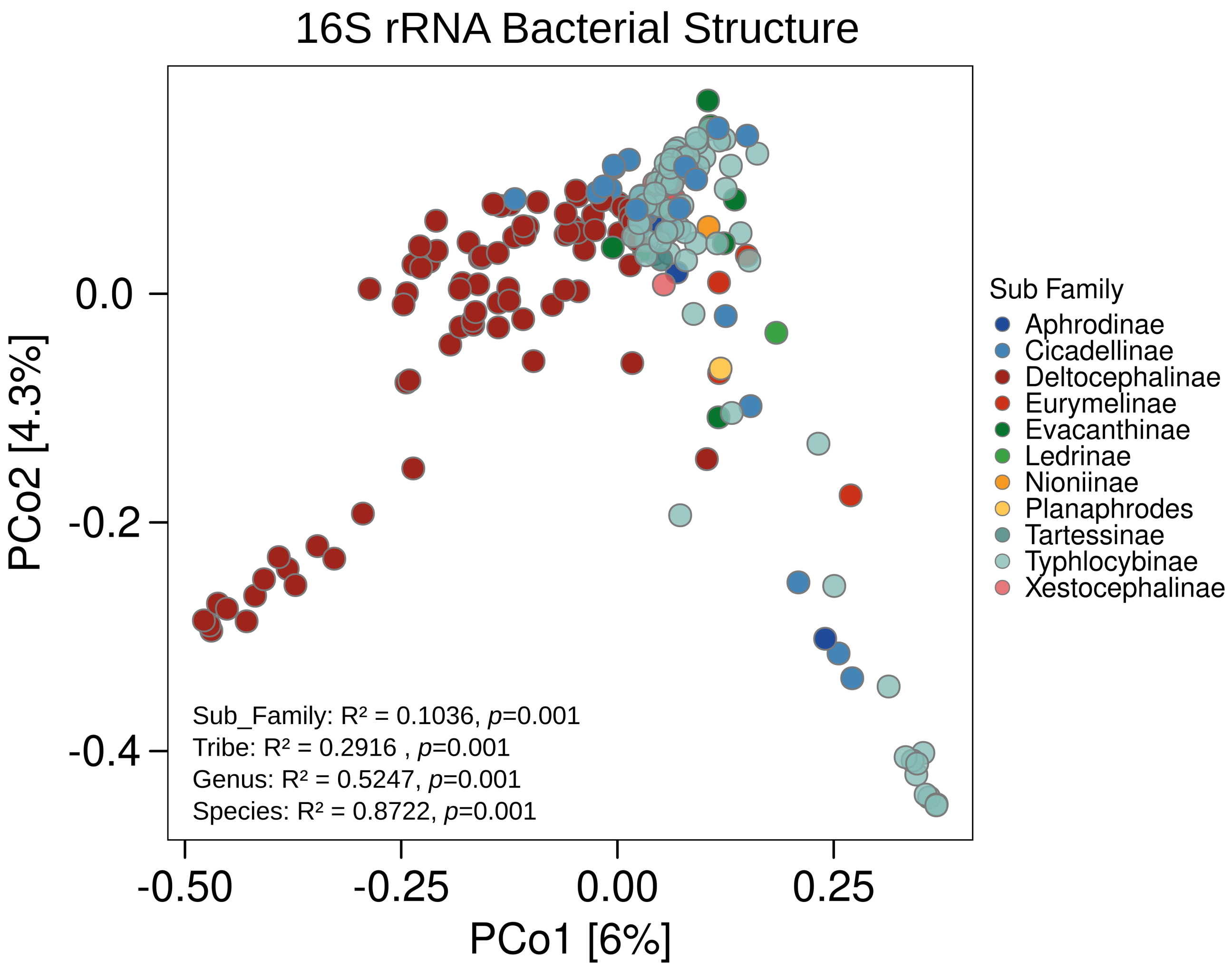


**Fig. S2 | Host phylogenetic signal in bacterial community structure based on 16S rRNA.** Principal Coordinate Analysis (PCoA) of 16S rRNA amplicon profiles from 264 leafhopper microbiomes colored by host subfamily. PERMANOVA statistics indicate that bacterial community composition exhibits increasing explanatory power at finer host taxonomic levels, with the strongest association observed at the species level. This supports the hypothesis that host evolutionary history partially constrains microbiome structure.

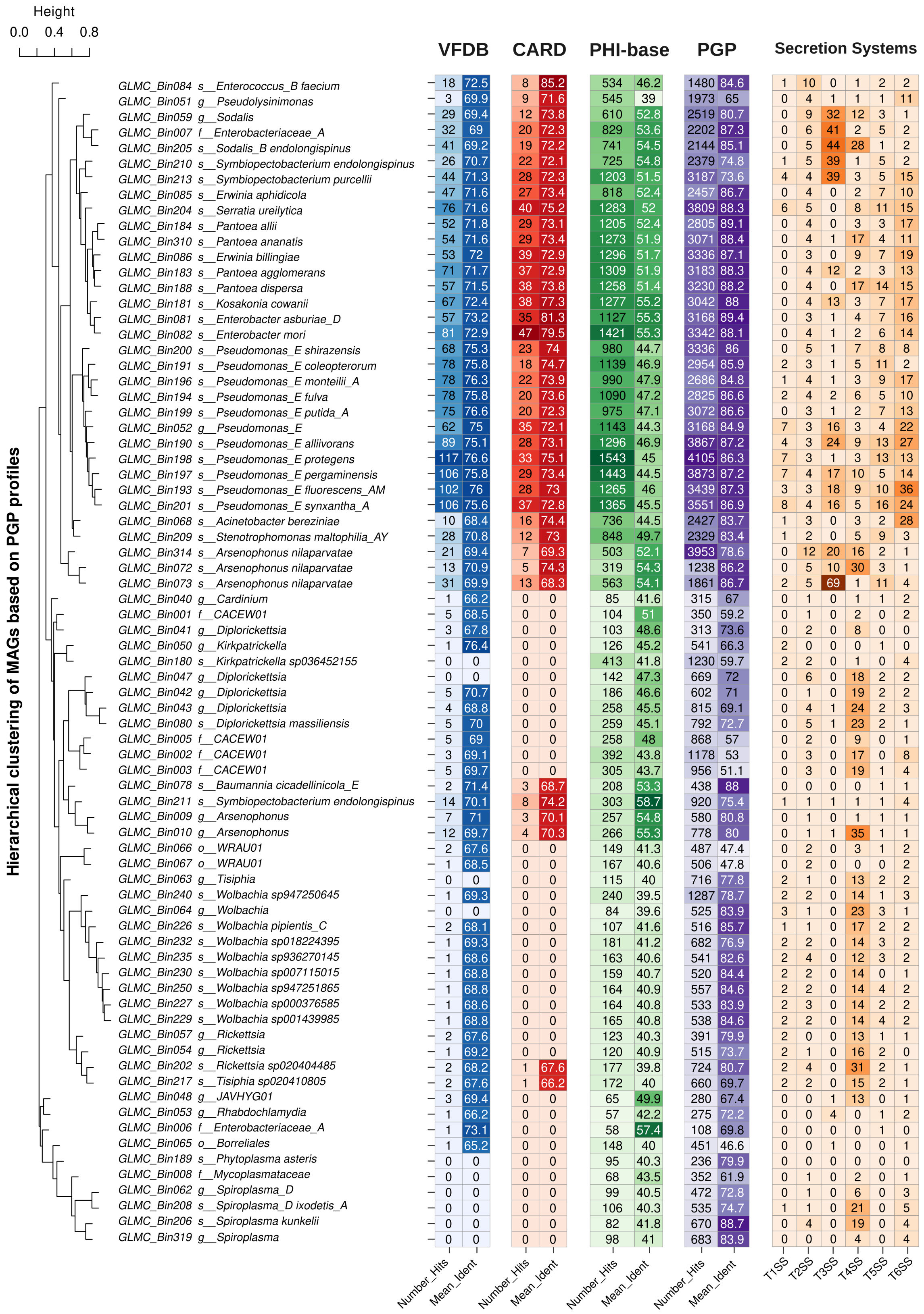
Height
0.0 0.4 0.8
Hierarchical clustering of MAGs based on PGP profiles
VFDB
CARD
PHI-base
PGP
Secretion Systems
Number_Hits Mean_Ident Number_Hits Mean_Ident Number_Hits Mean_Ident Number_Hits Mean_Ident T1SS T2SS T3SS T4SS T5SS T6SS
GLMC_Bin084 s__Enterococcus_B faecium 18 72.5 8 85.2 534 46.2 1480 84.6 1 10 0 1 2 2
GLMC_Bin051 g__Pseudolysinimonas 3 69.9 9 71.6 545 39 1973 65 0 4 1 1 1 11
GLMC_Bin059 g__Sodalis 29 69.4 12 73.8 610 52.8 2519 80.7 0 9 32 12 3 1
GLMC_Bin007 f__Enterobacteriaceae_A 32 69 20 72.3 829 53.6 2202 87.3 0 6 41 2 5 2
GLMC_Bin205 s__Sodalis_B endolongispinus 41 69.2 19 72.2 741 54.5 2144 85.1 0 5 44 28 1 2
GLMC_Bin210 s__Symbiopectobacterium endolongispinus 26 70.7 22 72.1 725 54.8 2379 74.8 1 5 39 1 5 2
GLMC_Bin213 s__Symbiopectobacterium purcellii 44 71.3 28 72.3 1203 51.5 3187 73.6 4 4 39 3 5 15
GLMC_Bin085 s__Erwinia aphidicola 47 71.6 27 73.4 818 52.4 2457 86.7 0 4 0 2 7 10
GLMC_Bin204 s__Serratia ureilytica 76 71.6 40 75.2 1283 52 3809 88.3 6 5 0 8 11 15
GLMC_Bin184 s__Pantoea allii 52 71.8 29 73.1 1205 52.4 2805 89.1 0 4 0 3 3 17
GLMC_Bin310 s__Pantoea ananatis 54 71.6 29 73.4 1273 51.9 3071 88.4 0 4 1 17 4 11
GLMC_Bin086 s__Erwinia billingiae 53 72 39 72.9 1296 51.7 3336 87.1 0 3 0 9 7 19
GLMC_Bin183 s__Pantoea agglomerans 71 71.7 37 72.9 1309 51.9 3183 88.3 0 4 12 2 3 13
GLMC_Bin188 s__Pantoea dispersa 57 71.5 38 73.8 1258 51.4 3230 88.2 0 4 0 17 14 15
GLMC_Bin181 s__Kosakonia cowanii 67 72.4 38 77.3 1277 55.2 3042 88 0 4 13 3 7 17
GLMC_Bin081 s__Enterobacter asburiae_D 57 73.2 35 81.3 1127 55.3 3168 89.4 0 3 1 4 7 16
GLMC_Bin082 s__Enterobacter mori 81 72.9 47 79.5 1421 55.3 3342 88.1 0 4 1 2 6 14
GLMC_Bin200 s__Pseudomonas_E shirazensis 68 75.3 23 74 980 44.7 3336 86 0 5 1 7 8 8
GLMC_Bin191 s__Pseudomonas_E coleopterorum 78 75.8 18 74.7 1139 46.9 2954 85.9 2 3 1 5 11 2
GLMC_Bin196 s__Pseudomonas_E monteilii_A 78 76.3 22 73.9 990 47.9 2686 84.8 1 4 1 3 9 17
GLMC_Bin194 s__Pseudomonas_E fulva 78 75.8 20 73.6 1090 47.2 2825 86.6 2 4 2 6 5 10
GLMC_Bin199 s__Pseudomonas_E putida_A 75 76.6 20 72.3 975 47.1 3072 86.6 0 3 1 2 7 13
GLMC_Bin052 g__Pseudomonas_E 62 75 35 72.1 1143 44.3 3168 84.9 7 3 16 3 4 22
GLMC_Bin190 s__Pseudomonas_E alliivorans 89 75.1 28 73.1 1296 46.9 3867 87.2 4 3 24 9 13 27
GLMC_Bin198 s__Pseudomonas_E protegens 117 76.6 33 75.1 1543 45 4105 86.3 7 3 1 3 13 13
GLMC_Bin197 s__Pseudomonas_E pergaminensis 106 75.8 29 73.4 1443 44.5 3873 87.2 7 4 17 10 5 14
GLMC_Bin193 s__Pseudomonas_E fluorescens_AM 102 76 28 73 1265 46 3439 87.3 3 3 18 9 10 36
GLMC_Bin201 s__Pseudomonas_E synxantha_A 106 75.6 37 72.8 1365 45.5 3551 86.9 8 4 16 5 16 24
GLMC_Bin068 s__Acinetobacter bereziniae 10 68.4 16 74.4 736 44.5 2427 83.7 1 3 0 3 2 28
GLMC_Bin209 s__Stenotrophomonas maltophilia_AY 28 70.8 12 73 848 49.7 2329 83.4 1 2 0 5 9 3
GLMC_Bin314 s__Arsenophonus nilaparvatae 21 69.4 7 69.3 503 52.1 3953 78.6 0 12 20 16 2 1
GLMC_Bin072 s__Arsenophonus nilaparvatae 13 70.9 5 74.3 319 54.3 1238 86.2 0 5 10 30 3 1
GLMC_Bin073 s__Arsenophonus nilaparvatae 31 69.9 13 68.3 563 54.1 1861 86.7 2 5 69 1 11 4
GLMC_Bin040 g__Cardinium 1 66.2 0 0 85 41.6 315 67 0 1 0 1 1 2
GLMC_Bin001 f__CACEW01 5 68.5 0 0 104 51 350 59.2 0 1 0 2 0 2
GLMC_Bin041 g__Diplorickettsia 3 67.8 0 0 103 48.6 313 73.6 0 2 0 8 0 0
GLMC_Bin050 g__Kirkpatrickella 1 76.4 0 0 126 45.2 541 66.3 2 0 0 0 1 0
GLMC_Bin180 s__Kirkpatrickella sp036452155 0 0 0 0 413 41.8 1230 59.7 2 2 0 1 0 4
GLMC_Bin047 g__Diplorickettsia 0 0 0 0 142 47.3 669 72 0 6 0 18 2 2
GLMC_Bin042 g__Diplorickettsia 5 70.7 0 0 186 46.6 602 71 0 1 0 19 2 2
GLMC_Bin043 g__Diplorickettsia 4 68.8 0 0 258 45.5 815 69.1 0 4 1 24 2 3
GLMC_Bin080 s__Diplorickettsia massiliensis 5 70 0 0 259 45.1 792 72.7 0 5 1 23 2 1
GLMC_Bin005 f__CACEW01 5 69 0 0 258 48 868 57 0 2 0 9 0 1
GLMC_Bin002 f__CACEW01 3 69.1 0 0 392 43.8 1178 53 0 3 0 17 0 8
GLMC_Bin003 f__CACEW01 5 69.7 0 0 305 43.7 956 51.1 0 3 0 19 1 4
GLMC_Bin078 s__Baumannia cicadellinicola_E 2 71.4 3 68.7 208 53.3 438 88 0 0 0 0 1 1
GLMC_Bin211 s__Symbiopectobacterium endolongispinus 14 70.1 8 74.2 303 58.7 920 75.4 1 1 1 1 1 4
GLMC_Bin009 g__Arsenophonus 7 71 3 70.1 257 54.8 580 80.8 0 1 0 1 1 1
GLMC_Bin010 g__Arsenophonus 12 69.7 4 70.3 266 55.3 778 80 0 1 1 35 1 1
GLMC_Bin066 o__WRAU01 2 67.6 0 0 149 41.3 487 47.4 0 2 0 3 1 2
GLMC_Bin067 o__WRAU01 1 68.5 0 0 167 40.6 506 47.8 0 2 0 0 0 2
GLMC_Bin063 g__Tisiphia 0 0 0 0 115 40 716 77.8 2 1 0 13 2 2
GLMC_Bin240 s__Wolbachia sp947250645 1 69.3 0 0 240 39.5 1287 78.7 2 2 0 14 1 3
GLMC_Bin064 g__Wolbachia 0 0 0 0 84 39.6 525 83.9 3 1 0 23 3 1
GLMC_Bin226 s__Wolbachia pipientis_C 2 68.1 0 0 107 41.6 516 85.7 1 1 0 17 2 2
GLMC_Bin232 s__Wolbachia sp018224395 1 69.3 0 0 181 41.2 682 76.9 2 2 0 14 3 2
GLMC_Bin235 s__Wolbachia sp936270145 1 68.6 0 0 163 40.6 541 82.6 2 4 0 12 3 2
GLMC_Bin230 s__Wolbachia sp007115015 1 68.8 0 0 159 40.7 520 84.4 2 2 0 14 0 1
GLMC_Bin250 s__Wolbachia sp947251865 1 68.8 0 0 164 40.9 557 84.6 2 2 0 14 4 2
GLMC_Bin227 s__Wolbachia sp000376585 1 68.6 0 0 164 40.8 533 83.9 2 3 0 14 2 2
GLMC_Bin229 s__Wolbachia sp001439985 1 68.8 0 0 165 40.8 538 84.6 2 2 0 14 4 2
GLMC_Bin057 g__Rickettsia 2 67.6 0 0 123 40.3 391 79.9 2 0 0 13 1 1
GLMC_Bin054 g__Rickettsia 1 69.2 0 0 120 40.9 515 73.7 2 1 0 16 2 0
GLMC_Bin202 s__Rickettsia sp020404485 2 68.2 1 67.6 177 39.8 724 80.7 2 4 0 31 2 1
GLMC_Bin217 s__Tisiphia sp020410805 2 67.6 1 66.2 172 40 660 69.7 2 2 0 15 2 1
GLMC_Bin048 g__JAVHYG01 3 69.4 0 0 65 49.9 280 67.4 0 0 1 13 0 1
GLMC_Bin053 g__Rhabdochlamydia 1 66.2 0 0 57 42.2 275 72.2 0 0 4 0 1 2
GLMC_Bin006 f__Enterobacteriaceae_A 1 73.1 0 0 58 57.4 108 69.8 0 0 0 0 1 0
GLMC_Bin065 o__Borreliales 1 65.2 0 0 148 40 451 46.6 0 0 1 0 0 1
GLMC_Bin189 s__Phytoplasma asteris 0 0 0 0 95 40.3 236 79.9 0 0 0 0 0 0
GLMC_Bin008 f__Mycoplasmataceae 0 0 0 0 68 43.5 352 61.9 0 1 0 2 0 1
GLMC_Bin062 g__Spiroplasma_D 0 0 0 0 99 40.5 472 72.8 0 1 0 6 0 3
GLMC_Bin208 s__Spiroplasma_D ixodetis_A 0 0 0 0 106 40.3 535 74.7 1 1 0 21 0 5
GLMC_Bin206 s__Spiroplasma kunkelii 0 0 0 0 82 41.8 670 88.7 0 4 0 19 0 4
GLMC_Bin319 g__Spiroplasma 0 0 0 0 98 41 683 83.9 0 0 0 4 0 4

**Fig. S3 | Functional profiling and hierarchical clustering of potentially pathogenic MAGs.** The dendrogram (left) groups Metagenome-Assembled Genomes (MAGs) based on PGP profiles, selecting for genera with known pathogenic members or suspected virulence potential. The heatmap displays annotation results across five categories: VFDB (Virulence Factor Database), CARD (Comprehensive Antibiotic Resistance Database), PHI-base (Pathogen-Host Interaction database), PGP (PLaBAse), and Secretion Systems (Type I–VI). For the first four categories, paired columns indicate the total number of gene hits (left sub-column) and the mean percentage identity (right sub-column). The Secretion Systems panel shows the count of identified components. Tile color intensity correlates with the magnitude of the values, which are explicitly annotated within each cell for clarity.

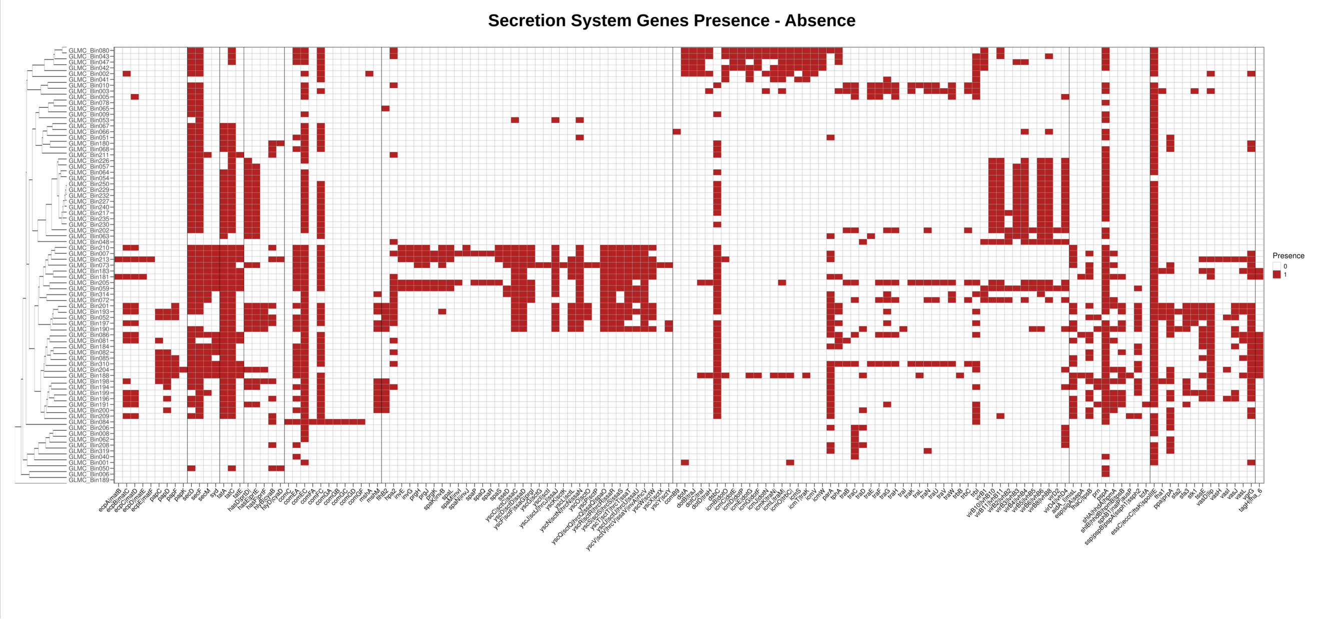


**Fig. S4 | Presence-absence profile of secretion system genes.** A binary heatmap displaying the specific gene repertoire for bacterial Secretion Systems I through VI across representative Metagenome-Assembled Genomes (MAGs) in the Global Leafhopper Microbiome Catalog (GLMC). The Y-axis lists the MAGs (arranged by hierarchical clustering), while the X-axis corresponds to individual genes encoding structural and effector components. Tile color indicates genomic conservation: red represents the presence of the specific gene, and white denotes its absence.

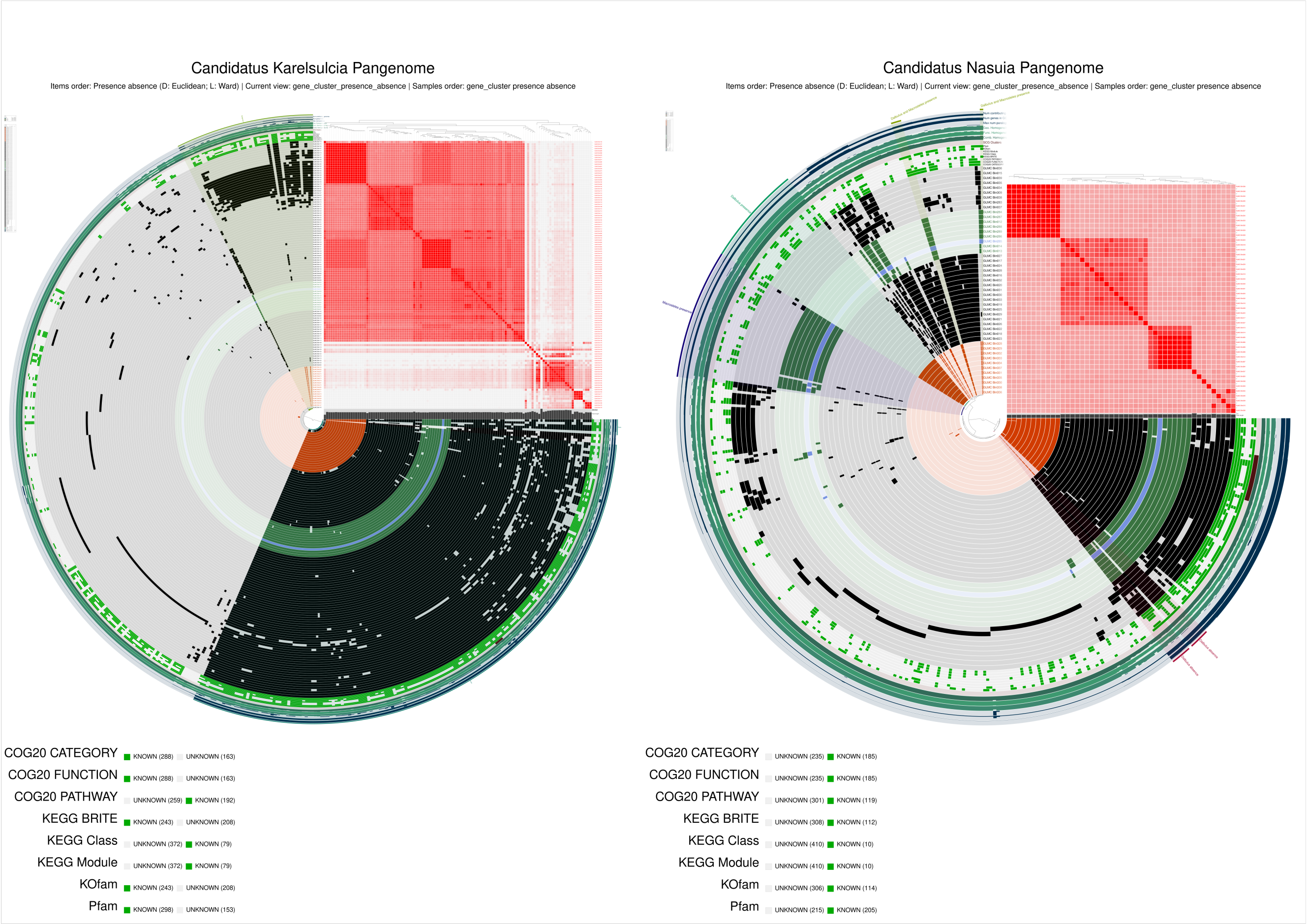


**Fig. S5 | Comparative pangenomics of the obligate symbionts '*Candidatus* Karelsulcia' and '*Ca.* Nasuia'.** Pangenome analysis of the two dominant obligate symbionts, (Left) '*Ca*. Karelsulcia' and (Right) '*Ca*. Nasuia'. In these circular plots, radial layers (rings) represent individual MAGs, and radial rays represent gene clusters. Dark/black segments indicate the presence of a gene cluster, while light/gap segments indicate absence. Genomes are organized by hierarchical clustering based on gene cluster distribution (central dendrogram). Inner metadata rings distinguish the host source: MAGs derived from *Macrosteles* species are colored green, while those from *Dalbulus* are orange. The inset heatmap (top right of each circle) displays pairwise Average Nucleotide Identity (ANI) values, highlighting genomic similarity alongside gene content. Outer layers summarize gene cluster statistics, including single-copy core genes (SCG), geometric and functional homogeneity, and maximum number of paralogs. Green distinct rings in the outer layers mark clusters with functional annotations in Pfam, KEGG, or COG databases. The comparison reveals that the '*Ca*. Karelsulcia' pangenome is significantly more stable and conserved than the structurally variable '*Ca*. Nasuia' pangenome.

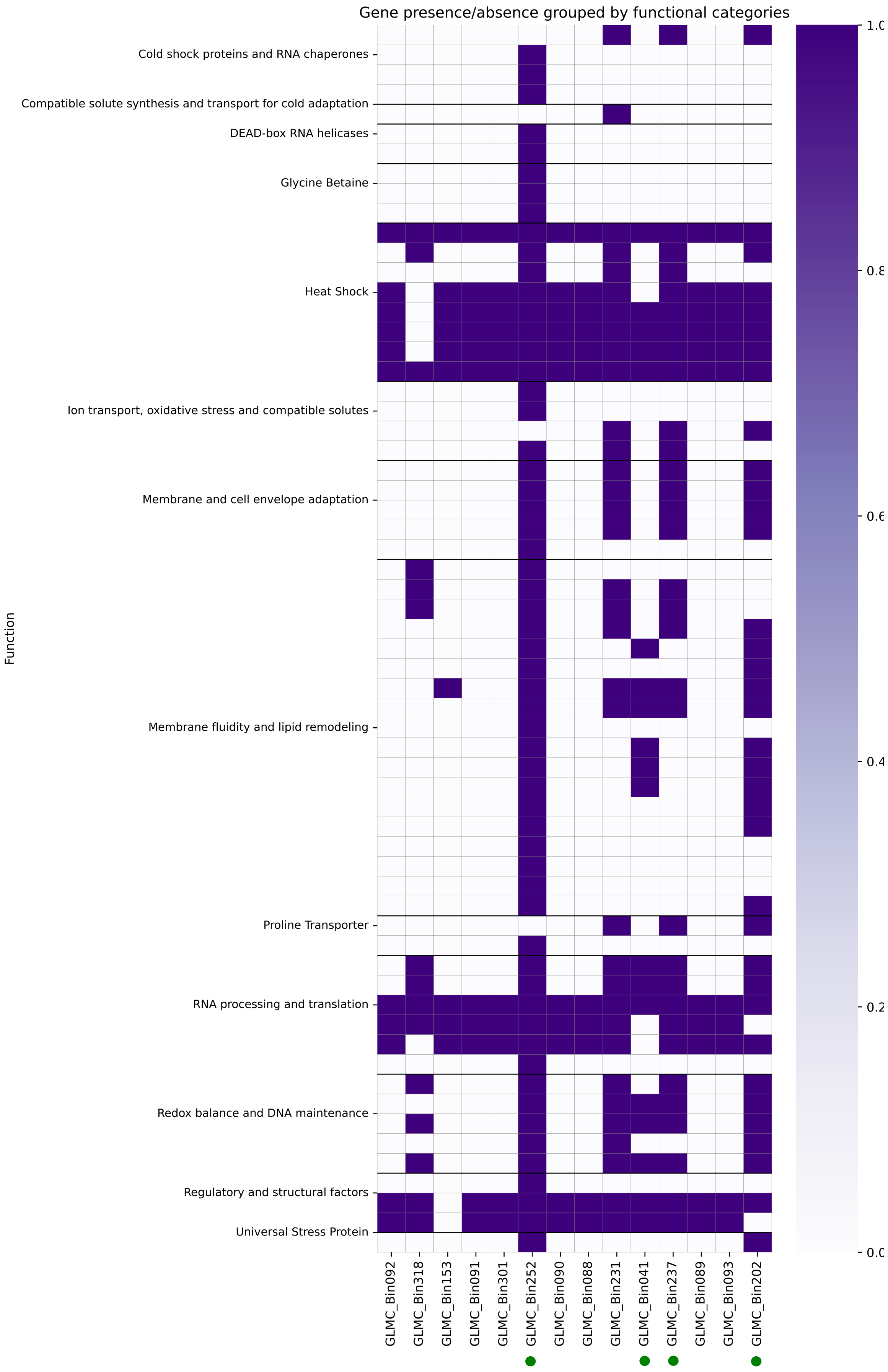
Gene presence/absence grouped by functional categories
Cold shock proteins and RNA chaperones
Compatible solute synthesis and transport for cold adaptation
DEAD-box RNA helicases
Glycine Betaine
Heat Shock
Ion transport, oxidative stress and compatible solutes
Membrane and cell envelope adaptation
Membrane fluidity and lipid remodeling
Proline Transporter
RNA processing and translation
Redox balance and DNA maintenance
Regulatory and structural factors
Universal Stress Protein
Function
GLMC_Bin092
GLMC_Bin318
GLMC_Bin153
GLMC_Bin091
GLMC_Bin301
GLMC_Bin252
GLMC_Bin090
GLMC_Bin088
GLMC_Bin231
GLMC_Bin041
GLMC_Bin237
GLMC_Bin089
GLMC_Bin093
GLMC_Bin202

**Fig. S6 | Distribution of abiotic stress tolerance genes in Arctic leafhopper microbiomes.** Binary heatmap profiling the presence of genes conferring resistance to environmental stressors in Metagenome-Assembled Genomes (MAGs) recovered from the Canadian Arctic (Nunavik). Rows correspond to specific stress response genes, while columns represent individual MAGs. Purple tiles indicate the presence of a gene, and white tiles denote its absence. Green circles highlight MAGs derived from *Macrosteles* host populations as well as the *Arsenophonus* lineage, distinguishing them from other recovered taxa.

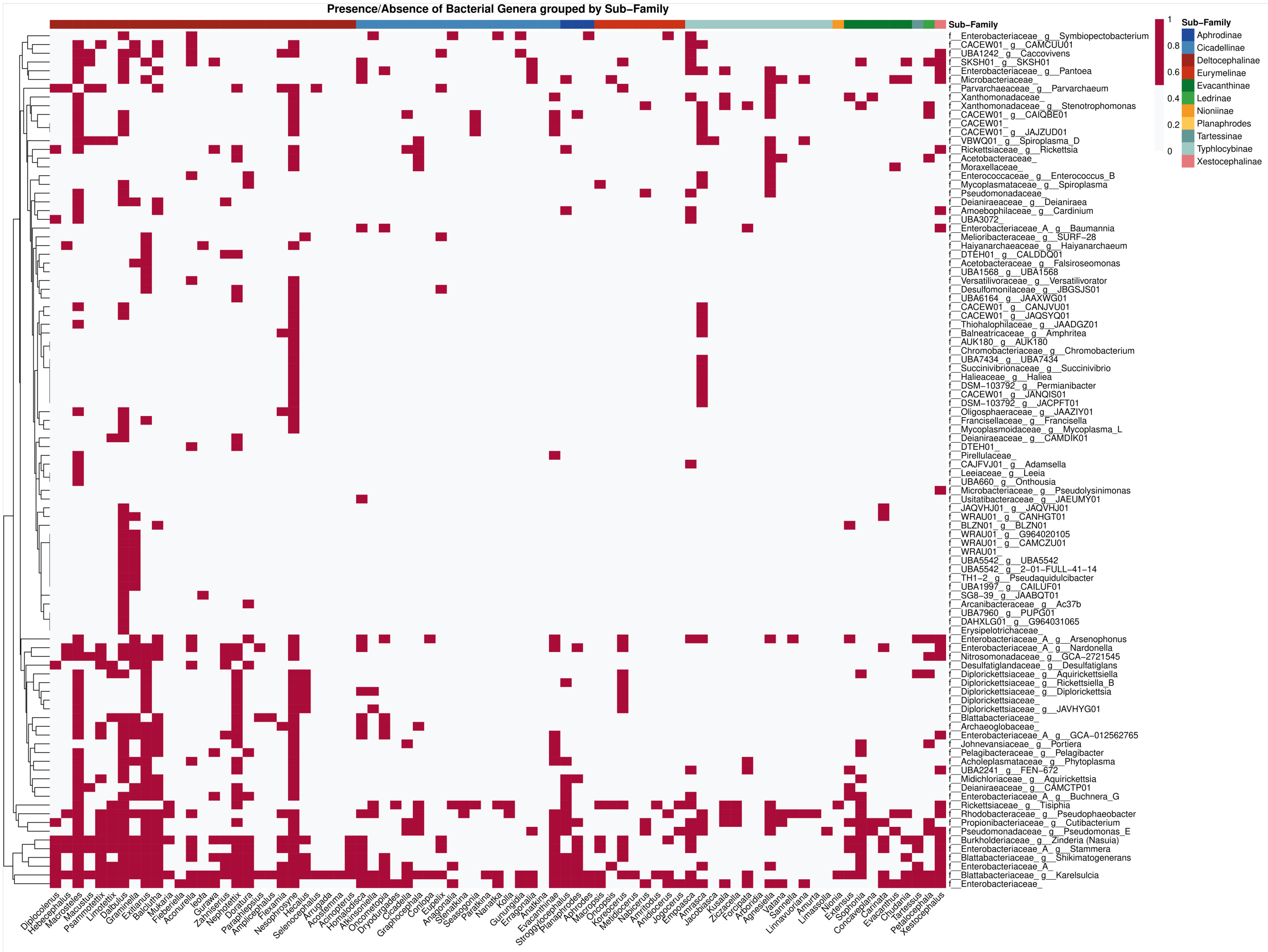


**Fig. S7 | Prokaryotic taxonomic profiling across leafhopper lineages.** A binary heatmap displaying the distribution of bacterial genera identified via SingleM profiling using the GTDB marker gene database. Rows represent bacterial taxonomy (clustered hierarchically), while columns represent leafhopper host genera, grouped by their respective subfamilies (indicated by the top color bar). Red tiles indicate the presence of a bacterial genus, and white tiles denote its absence. The dense cluster at the bottom highlights the widespread prevalence of obligate symbionts, including ‘*Candidatus* Karelsulcia’, ‘*Ca*. Nasuia’, and ‘*Ca*. Baumannia’, across diverse host lineages.

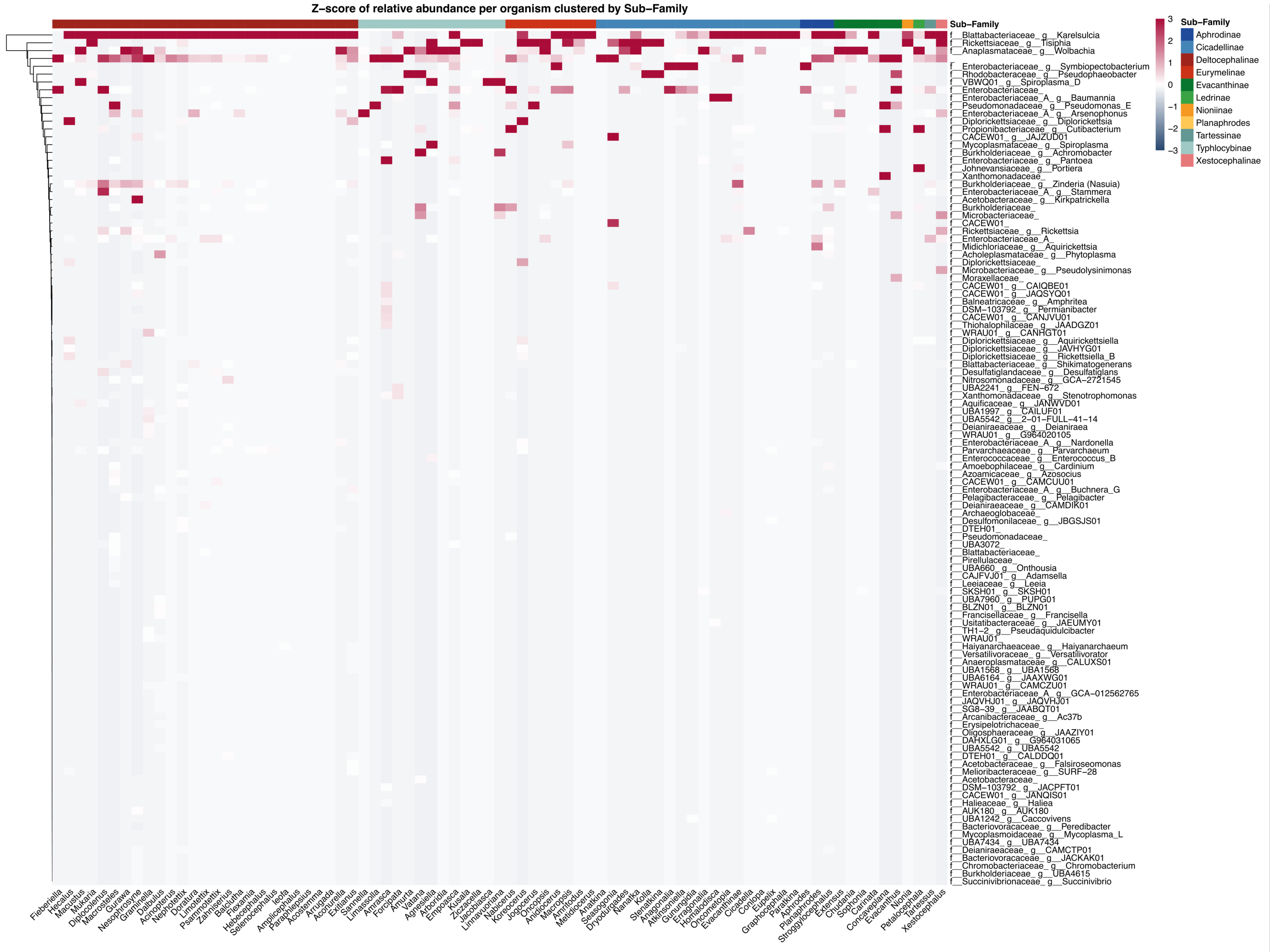


**Fig. S8 | Prokaryotic taxonomic profiling across leafhopper lineages.** Heatmap displaying the standardized relative abundance (Z-scores) of bacterial genera identified via SingleM profiling (GTDB). Rows represent hierarchically clustered bacterial taxa, while columns represent leafhopper host genera grouped by subfamily (indicated by the top color bar). Tile color intensity reflects the Z-score, where deeper red indicates higher relative abundance compared to the row mean. The analysis reveals a distinct high-abundance cluster at the bottom comprising obligate symbionts ‘*Candidatus* Karelsulcia’, ‘*Ca.* Nasuia’, and ‘*Ca.* Baumannia’, which contrasts with the upper or more variable abundance of facultative and environmental taxa in the lower clusters.
